# Computational Circuit Mechanisms Underlying Thalamic Control of Attention

**DOI:** 10.1101/2020.09.16.300749

**Authors:** Qinglong L. Gu, Norman H. Lam, Ralf D. Wimmer, Michael M. Halassa, John D. Murray

## Abstract

The thalamus engages in attention by amplifying relevant signals and filtering distractors. Whether architectural features of thalamic circuitry offer a unique locus for attentional control is unknown. We developed a circuit model of excitatory thalamocortical and inhibitory reticular neurons, capturing key observations from task-engaged animals. We found that top-down inputs onto reticular neurons regulate thalamic gain effectively, compared to direct thalamocortical inputs. This mechanism enhances downstream readout, improving detection, discrimination, and cross-modal performance. The model revealed heterogeneous thalamic responses that enable decoding top-down versus bottom-up signals. Spiking activity from task-performing mice supported model predictions, with a similar coding geometry in auditory thalamus and readout strategy in auditory cortex. Dynamical systems analysis explained why reticular neurons are potent sites for control, and how lack of excitatory connectivity among thalamocortical neurons enables separation of top-down from bottom-up signals. Our work reveals mechanisms for attentional control and connects circuit architectures to computational functions.

## Introduction

Attention is a core cognitive function that involves flexible prioritization of sensory inputs. In an environment with multiple stimuli or features, attentional modulation can enhance processing of selected sensory signals and filter out distractors in a goal-directed manner. A growing experimental literature shows that selective attention engages subcortical structures (Krauzlis et al., 2018), including the thalamus (Saalmann and Kastner, 2011; Halassa and Kastner, 2017). Within sensory thalamic structures, glutamatergic thalamocortical (TC) neurons process and transmit information to cortex for downstream computation. The thalamic reticular nucleus (TRN), which forms a thin capsule around thalamic nuclei, contains GABAergic reticular (RE) neurons that provide inhibition onto TC neurons (Jones, 2012). In an influential theoretical proposal, Crick (1984) hypothesized that selective attention could be implemented with the TRN acting as an attentional “spotlight,” suggesting that “if the thalamus is the gateway to the cortex, the reticular complex might be described as the guardian of the gateway.”

Studies in rodents, monkeys, and humans characterize sensory thalamus as a locus of attentional filtering under flexible top-down control via the TRN (O’Connor et al., 2002; McAlonan et al., 2008; Ahrens et al., 2015; Wimmer et al., 2015; Nakajima et al., 2019). Wimmer et al. (2015) recorded neuronal spiking activity from the visual TRN and lateral geniculate nucleus (LGN) in mice trained on a cross-modal attention task, in which a cue informed of the subject of which sensory modality (vision vs. audition) to attend to. TRN and LGN exhibited opposite modulations by attentional selection: when vision was the attended modality, visual TRN decreased firing activity and LGN exhibited increased activity and higher gain of stimulus response. Nakajima et al. (2019) found a pathway for this attentional control of thalamus, from the prefrontal cortex through the basal ganglia’s inhibitory projections onto the TRN.

These convergent empirical findings raise key computational questions about the mechanisms and functions of attentional control in thalamic circuits. At the mechanistic level of circuit computations, why the thalamus is as an effective locus for attentional control, and why control via the TRN, compared to compared to direct projections to TC neurons, may be more effective for regulation of thalamic processing are open questions. Furthermore, how downstream readout of thalamic activity in cortex can parse bottom-up from top-down signals are also unknown. Computational modeling provides a useful framework for understanding how neural circuit mechanisms can subserve cognitive functions. Thalamic dynamics such as oscillations have been modeled at the neuronal and microcircuit levels (Destexhe et al., 1993b; 1994; 1996; Wang et al., 1995; Bazhenov et al., 2000; Sohal et al., 2006). However, there has been relatively little biophysically grounded modeling of thalamic circuits in dynamical regimes consistent with neural recordings from alert behaving animals, which could yield insight into the interplay of cellular and circuit mechanisms supporting attentional modulation.

To investigate these issues, we developed a biophysically grounded thalamic circuit model and studied how modulation by top-down inputs can implement attentional control of stimulus processing. We found that this model, with circuit dynamics constrained by experiments to operate in the regime of alert behaving animals, can capture key experimental observations of thalamic activity and its modulation by attention. Dynamical systems analysis explains why RE neurons are an especially potent site for top-down control, compared to TC neurons. The geometry of heterogeneous thalamic responses makes predictions for downstream readout of information from the confluence of bottom-up and top-down signals, for attentional enhancement of performance in detection, discrimination, and cross-modal task paradigms. Motivated by model predictions, we examined spiking activity from auditory thalamus (ventral medial geniculate body [MGBv]) during the two-alternative forced choice task featuring modality-selective attention (Nakajima et al., 2019), and found the existence of the similar coding geometrical structure. Analysis of spiking activity in primary auditory cortex (A1) during this task further supports that thalamic gain modulation arise from local TC-RE circuit computations which can be effectively read out by cortical circuits. Our sensitivity analysis reveals how dynamical and computational features depend on excitation-inhibition balance. Finally, modeling shows that strong local recurrent excitation can impair downstream decoding of multiplexed bottom-up and top-down signals, which suggests why thalamic circuits, lacking such recurrent excitation, is an effective locus for attentional modulation. The model therefore provides theoretical insight into circuit mechanisms of attentional control, parsimoniously captures a range of experimental observations, and makes experimentally testable predictions for future studies of the thalamus’s role in cognitive function.

## Results

### Thalamic Circuit Architecture

In this study we developed a thalamic circuit model of attention which is parsimonious yet grounded in physiology and anatomy of the system (**Figure 1**; see **Methods**). The model consists of a population of excitatory thalamocortical (TC) neurons, and another of inhibitory reticular (RE) neurons. TC neurons receive stimulus inputs, from a particular modality, and TC outputs are considered to project to downstream areas for readout and processing. RE neurons are reciprocally connected to TC neurons but also receive external inputs, including top-down attentional signals. Various forms of top-down inputs, mediating modality-selective attentional control signals, may also target TC neurons to modulate circuit responses.

**Figure 1:**
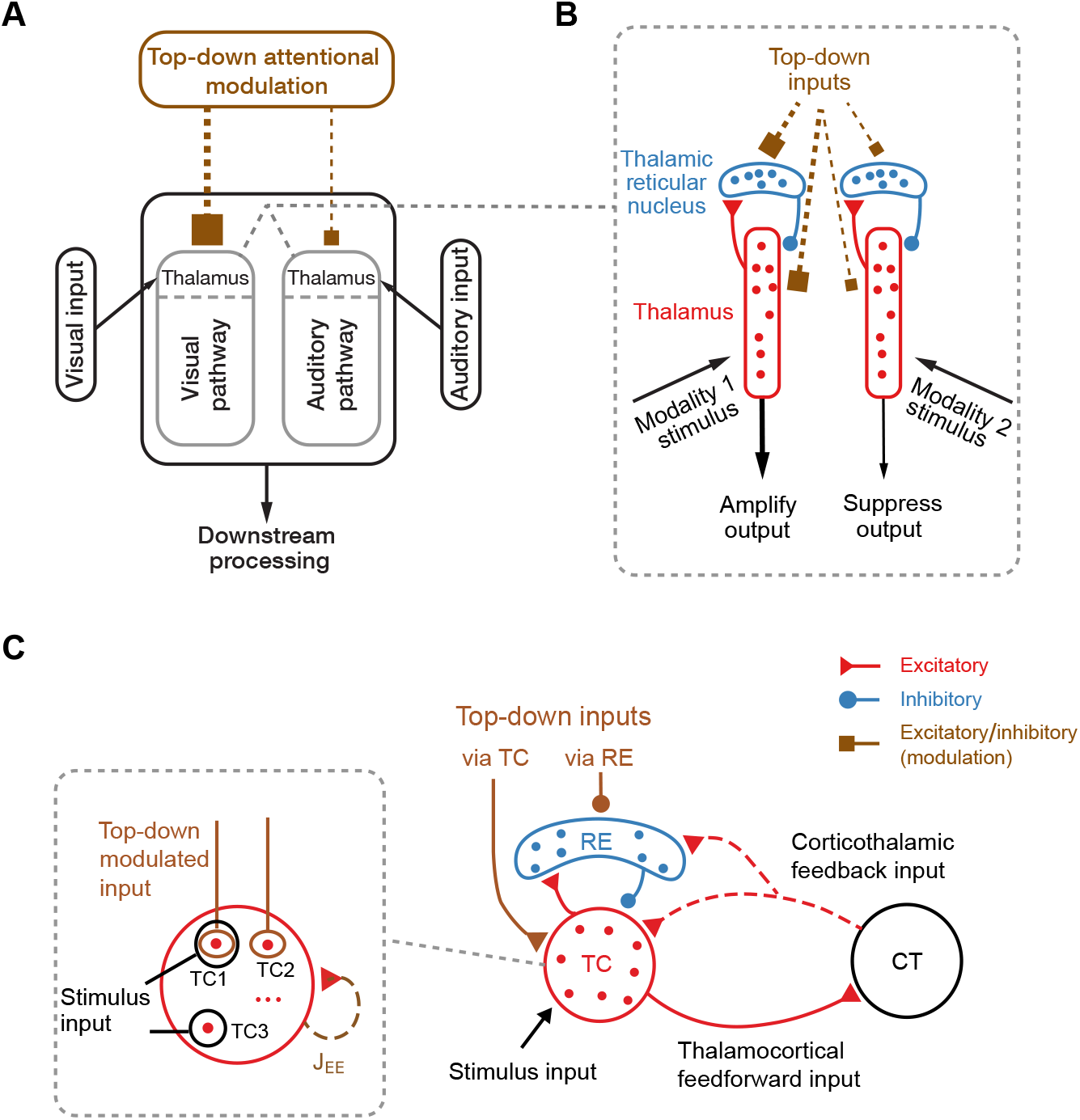
Model architecture for attentional control of thalamic circuits. **(A)** Conceptual schematic of modality-selective attention. When receiving simultaneous stimulus inputs from multiple modalities (e.g., visual and auditory) for processing in downstream cortical areas, top-down inputs onto thalamic structures can mediate modality-selective attentional modulation of thalamic responses. **(B)** Thalamic circuit implementation of top-down modulation on selective attention. Distinct populations of excitatory thalamocortical neurons (TC, red) receive stimulus inputs from separate modalities and project their outputs to the cortex. Each population of TC neurons is reciprocally coupled to an associated population of inhibitory reticular neurons (RE, blue). To implement attentional control, top-down inputs to RE or TC neurons modulate thalamic outputs to the cortex, e.g., through amplification of stimulus signals from an attended sensory modality or suppression of those from a distractor modality. **(C)** Circuit mechanisms for top-down attentional modulation. Major topics of study here include roles of heterogeneous of population coding for bottom-up and top-down signals in thalamus; differential potency of RE vs. TC cells as sites for top-down inputs; and inter-regional communication between thalamus and downstream cortical circuits.

To enable modeling of thalamic circuit dynamics during attentional control, we first constrained our circuit model to operate in a dynamical regime consistent with *in vivo* observations from alert behaving animals. Prior computational modeling studies of TC and RE neurons primarily focused on neuronal dynamics observed *in vitro*, under anesthesia, or during sleep, in which TC and RE neurons can intrinsically generate strong oscillations between bursting and hyperpolarization (Wang et al., 1995; Destexhe et al., 1996; 1998; Bazhenov et al., 2002; Krishnan et al., 2016; Timofeev et al., 2020). In contrast, spontaneous activity in the alert awake state is characterized by irregular, quasi-asynchronous spiking with only intermittent bursting and lack of strong intrinsic oscillation (Bruno, 2006; Temereanca et al., 2008; Ramcharan et al., 2005; Hirai et al., 2017; Bastos et al., 2014; Saleem et al., 2017). In addition, a strong and brief optogenetic depolarization of RE neurons evokes a transient response involving increased probability of TC bursting and an oscillatory bout in the spindle (7–15 Hz) frequency range (Halassa et al., 2011). Therefore, quasi-asynchronous spontaneous and evoked oscillatory activity can characterize the alert awake dynamical regime of a thalamic circuit and constrain model parameters (**Figure S1**).

Using the circuit model in this operating regime, we sought to answer the following major questions: What are functional differences between RE neurons and TC neurons as sites for top-down control? Can thalamic coding allow for attentional gain control yet enable downstream decoding to distinguish bottom-up stimulus and top-down control? Is thalamic gain modulation computed locally or inherited from cortical feedback? How would the architectural feature of thalamic lack of recurrent excitation influence its engagement in attentional control? (**Figure 1C**).

### Top-down Regulation of Thalamic Dynamics and Stimulus Processing

First, we sought to understand the effects of top-down control inputs and how they can implement attentional modulation of stimulus processing. We first consider our thalamic circuit of one modality with top-down inputs taking the form of inhibition onto the RE neurons of the TRN, based on the basal ganglia pathway described by Nakajima et al. (2019) (**Figure 2A**). We found that increasing the strength of top-down drive suppresses RE firing rates, and elevates TC firing rates via disinhibition, which captures experimental observations when attention is directed toward the associated sensory modality (Wimmer et al., 2015; Nakajima et al., 2019) (**Figure 2A**).

**Figure 2:**
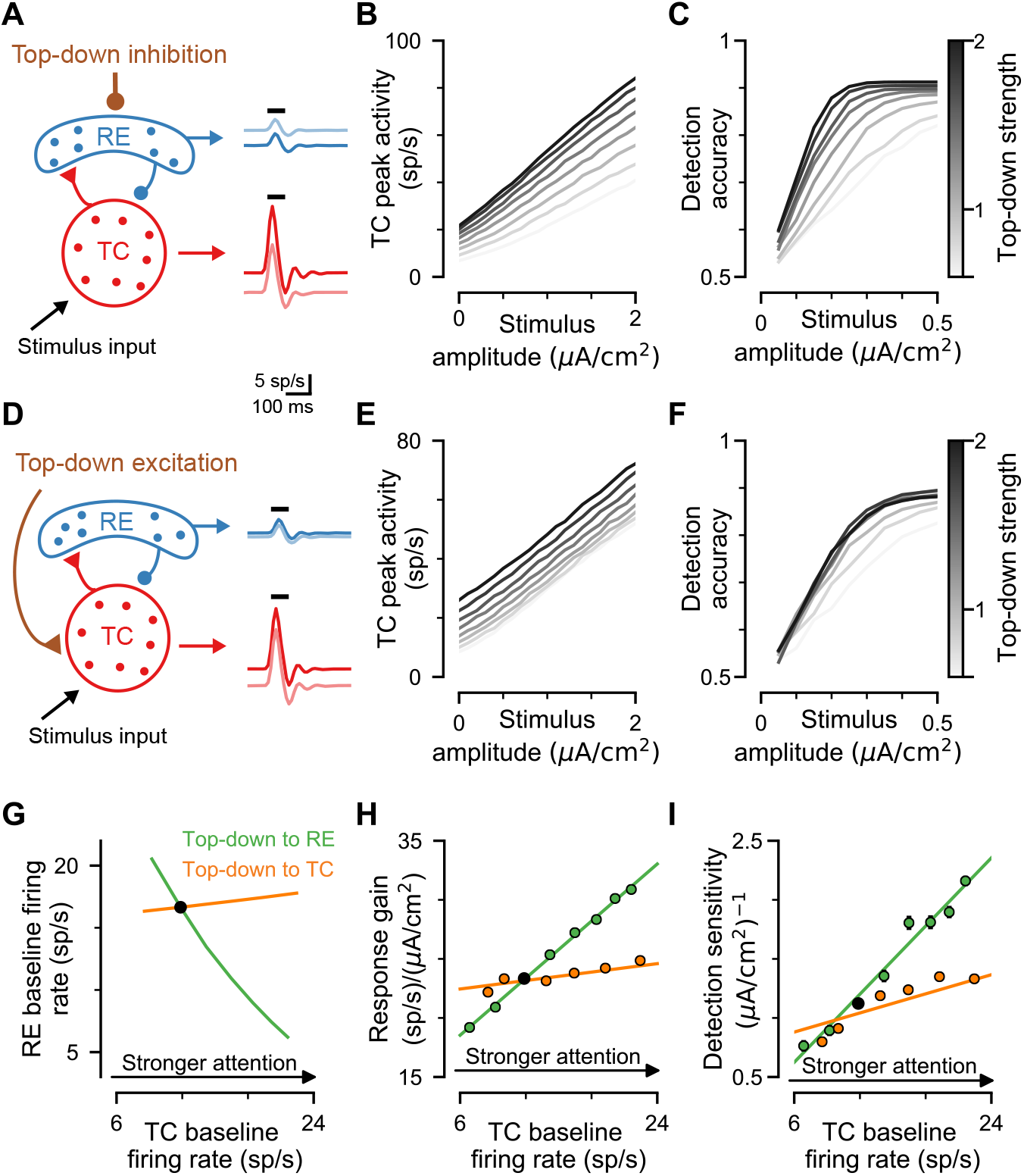
Top-down modulation via RE neurons, in contrast to TC neurons. **(A)** Schematic of top-down inhibition to RE neurons modulating thalamic response to stimulus input, with different top-down modulation strengths (dark for strong, light for weak). **(B)** Circuit response function, i.e. TC firing rate as a function of stimulus strength, under different strengths of top-down modulation. **(C)** Detection accuracy of a downstream decoder, which is based on a linear support vector machine (SVM) and operates on thalamic output firing-rate patterns. **(D–F)** Same as **(A–C)**, but with top-down modulation via excitation to TC neurons instead of RE. **(G)** RE baseline activity as a function of top-down modulation strength, for inhibition to RE or excitation to TC. To facilitate comparison between RE vs. TC modulations, top-down modulation strength is parameterized in terms of its net effect on TC baseline firing rate. Black dot denotes the control circuit. **(H)** Response gain as a function of top-down modulation strength. Response gain is defined as the slope of the TC response as a function of stimulus amplitude (from **B** and **E**). **(I)** Sensitivity of detection accuracy as a function of top-down modulation strength. Sensitivity is defined as the slope of the initial part of the accuracy curve (from **C** and **F**). Top-down modulation via RE neurons has a larger effect on gain and sensitivity than via TC neurons.

In addition to its effects on baseline activity, top-down inhibition of RE boosted the TC response to bottom-up stimulus inputs (**Figure 2B**). The TC response is a quasi-linear function of stimulus input amplitude, and strengthening the top-down input increases the gain of TC response (i.e., its slope as a function of stimulus amplitude). This increase in thalamic response gain by attention is also in line with experimental observations (Wimmer et al., 2015; Nakajima et al., 2019; McAlonan et al., 2008; O’Connor et al., 2002). Can this increase in TC response gain enhance downstream readout of the stimulus, for attentional improvement in behavior? To examine such questions, we built a decoder for stimulus detection, using the linear support vector machine (SVM) classifier framework, which is trained on the firing levels across the TC population (see **Methods**). We found that top-down modulation greatly improves detection accuracy through greater sensitivity (**Figure 2C**). These findings link top-down modulation of RE to increased TC response gain and improved downstream readout for psychophysical behavior.

### Reticular vs. Thalamocortical Neurons as Sites for Top-Down Control

Are RE and TC populations similarly potent sites for thalamic regulation by top-down inputs? As an alternative to disinhibition of TC via RE neurons, which indirectly modulates TC response gain as shown above, top-down excitatory inputs (e.g., from cortex) could directly project to TC neurons to modulate their activity (Briggs and Usrey, 2008; Béhuret et al., 2015). We found that elevating top-down excitation to TC neurons can strongly increase the baseline firing rate of TC neurons (**Figure 2D**). RE neurons exhibited a weak increase in baseline firing rate, in contrast to the strong decrease induced by top-down inhibition to RE (**Figure 2A, D and G**). RE vs. TC as candidate sites for top-down input are therefore dissociable in their effects on baseline RE activity (**Figure 2G**), and RE neurons as the primary site is consistent with experimental findings from Wimmer et al. (2015) and Nakajima et al. (2019).

Remarkably, top-down input to TC neurons exerted only a very weak effect on response gain, compared to top-down input to RE neurons (**Figure 2B, E and H**). Top-down input to TC neurons had a correspondingly weak effect on detection accuracy from the downstream SVM decoder (**Figure 2C, F and I**). Response gain and detection sensitivity increase monotonically with top-down modulation strength (**Figures 2H-I** and **S2**). Response gain and detection sensitivity were both much more potentially regulated by top-down input to RE neurons than top-down input to TC neurons (by factors of 6.8 and 3.6, respectively).

We also investigated a second canonical task paradigm, categorical discrimination, wherein the subject must distinguish which of two stimuli is presented. To study this paradigm, we introduced two stimulus patterns as inputs to the circuit, defined as two Gaussian-tuned inputs which partially overlap in their projections onto TC neurons (**Figure S3A**). Here, the top-down input onto the RE population is non-specific and unstructured, and therefore is not informed by the stimuli or discrimination boundary. To characterize the impact of top-down attentional modulation on downstream readout we trained an SVM decoder for stimulus classification (see **Methods**). Overall, we found that top-down modulation has a similar effect on discrimination as on detection. Top-down modulation via RE improved discrimination performance (**Figure S3B**), and was more effective, compared to TC, at enhancing discrimination sensitivity (**Figure S3C**).

### Dynamical Systems Analysis of Reduced Mean-Field Model

The above results reveal that RE neurons are more potent sites for top-down control of response gain, relative to TC neurons, with corresponding impact on downstream readout of stimulus signals. To understand the dynamical circuit mechanisms underlying this differential sensitivity to RE vs. TC modulation, we developed and analyzed a reduced mean-field model of the thalamic circuit. This reduced model comprises two dynamical variables representing the aggregate activity levels of the RE and and TC populations. The response gain of the circuit is an emergent property that is shaped by the interplay between the neuronal sensitivities and the feedback inhibition loop created by TC-RE interconnections (**Figure 3A**). Each neural population is governed by a neuronal transfer function describing the output firing rate as a function of the input current (commonly called an f-I curve), whose shape follows an expansive nonlinearity (**Figure 3B**). Neuronal and synaptic parameters were fit to, or derived from, the spiking circuit model (see **Methods**).

**Figure 3:**
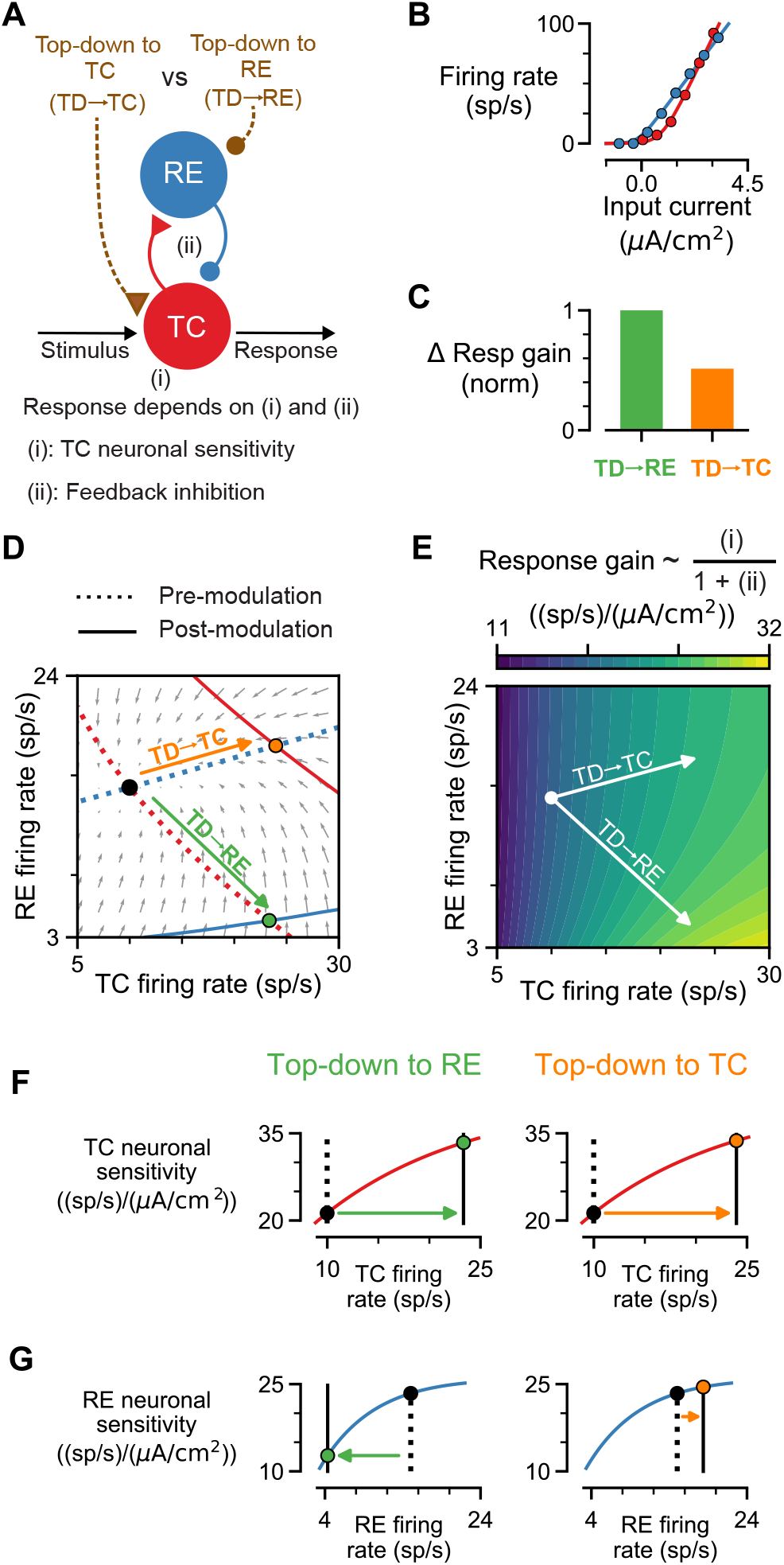
Mean-field analysis of gain control by top-down inputs. **(A)** Schematic of mean-field firing-rate circuit model, with TC and RE activity as two dynamical variables. Top-down (TD) inhibition to RE (TD→RE) or excitation to TC (TD→TC) modulates the circuit state. **(B)** Neuronal input-output transfer functions. Parameters were fit to simulated spiking data from the TC and RE single-neuron models (dots). **(C)** Efects of TD→RE and TD→TC modulation on circuit response gain. TD→RE vs. TD→TC pathways are compared by setting top-down strengths to have the same TC baseline firing rate (as shown in **D**). For ease of comparison, changes in response gain are normalized by that induced by TD→RE. TD→RE increases response gain more than TD→TC. **(D)** Phase-plane analysis showing nullclines of TC (red) and RE (blue) firing rates. Solid lines represent nullclines under top-down modulation. Intersections of nullclines are steady-state fixed points of the circuit (black for control, green for TD→RE, and orange for TD→TC). The small grey arrows represent the vector field for activity flow in the control state. **(E)** Circuit response gain as a function of TC and RE firing rates. **(F)** TC neuronal sensitivity, i.e. the slope of the TC neuronal transfer function, as a function of TC firing rate under top-down modulation via RE (left) or TC (right). Both top-down modulations can increase TC activity and its neuronal sensitivity at the similar level. **(G)** RE neuronal sensitivity as a function of firing rate, under top-down inputs to RE (left) or TC (right). TD→RE decreases RE firing rate (left) whereas TD→TC increases RE firing rate (right), resulting in distinct gain effects. Top-down inputs drive the circuit state from the black dot to colored dots.

We found that the reduced mean-field model exhibits key properties of the spiking circuit, including that RE neurons are a more potent site for top-down regulation of response gain, compared to TC neurons (by a factor of nearly two, **Figure 3C**). The steady-state TC and RE activity levels under specific inputs into the circuit. As in the spiking circuit, top-down input via TC vs. RE neurons produces dissociable changes in the firing rates of RE neurons, which reflects the non-inhibition-stabilized regime (Ozeki et al., 2009; Litwin-Kumar et al., 2016) (**Figure 3D**).

The analytical tractability of the reduced model allows calculation of the circuit response gain as a function of TC and RE activity levels. Top-down input onto RE, compared to TC, shifts the circuit to a state of higher response gain (**Figure 3E**). The mechanistic basis of this difference is revealed by the analytical expression for response gain (**Equation 16** in **Methods**) with dependence on the RE and TC neuronal sensitivities, i.e., the slopes of the neuronal input-output transfer functions (**Figure 3B**): response gain increases with higher TC neuronal sensitivity (and firing rate), and decreases with higher RE neuronal sensitivity (and firing rate). TC neuronal sensitivity is increased by top-down input to RE or to TC (**Figure 3F**). With top-down inhibition to RE, RE neuronal sensitivity decreases (**Figure 3G** left), which in turn boosts response gain. In contrast, with top-down excitation to TC, RE neuronal sensitivity increases (**Figure 3G** right), which in turn lowers response gain and counteracts the contribution from TC sensitivity. These analyses thereby give insight into how the interplay of neuronal and circuit properties give rise to the differential potency of RE as a site for top-down control of thalamic gain.

### Downstream Decoding via Heterogeneous Population Activity

As shown above in the spiking circuit, top-down modulation via RE neurons improves performance of a downstream decoder trained for stimulus detection (**Figure 2C, I**). We next investigated the population coding principles by which downstream readout can be facilitated by top-down modulation of distributed TC response patterns. The decoder we implemented is a linear support vector machine (SVM) trained on the patterns of firing-rate activity across the heterogeneous population of 1000 TC neurons (**Figure 4A**) (see **Methods**). The population-activity decoder can be viewed geometrically as a classifier hyperplane in the high-dimensional space of neuronal activity (**Figure 4B**). To examine whether a distributed code across heterogeneous neurons is important for decoding, we also considered a second decoder whose input is only the mean firing rate averaged across the TC population.

**Figure 4:**
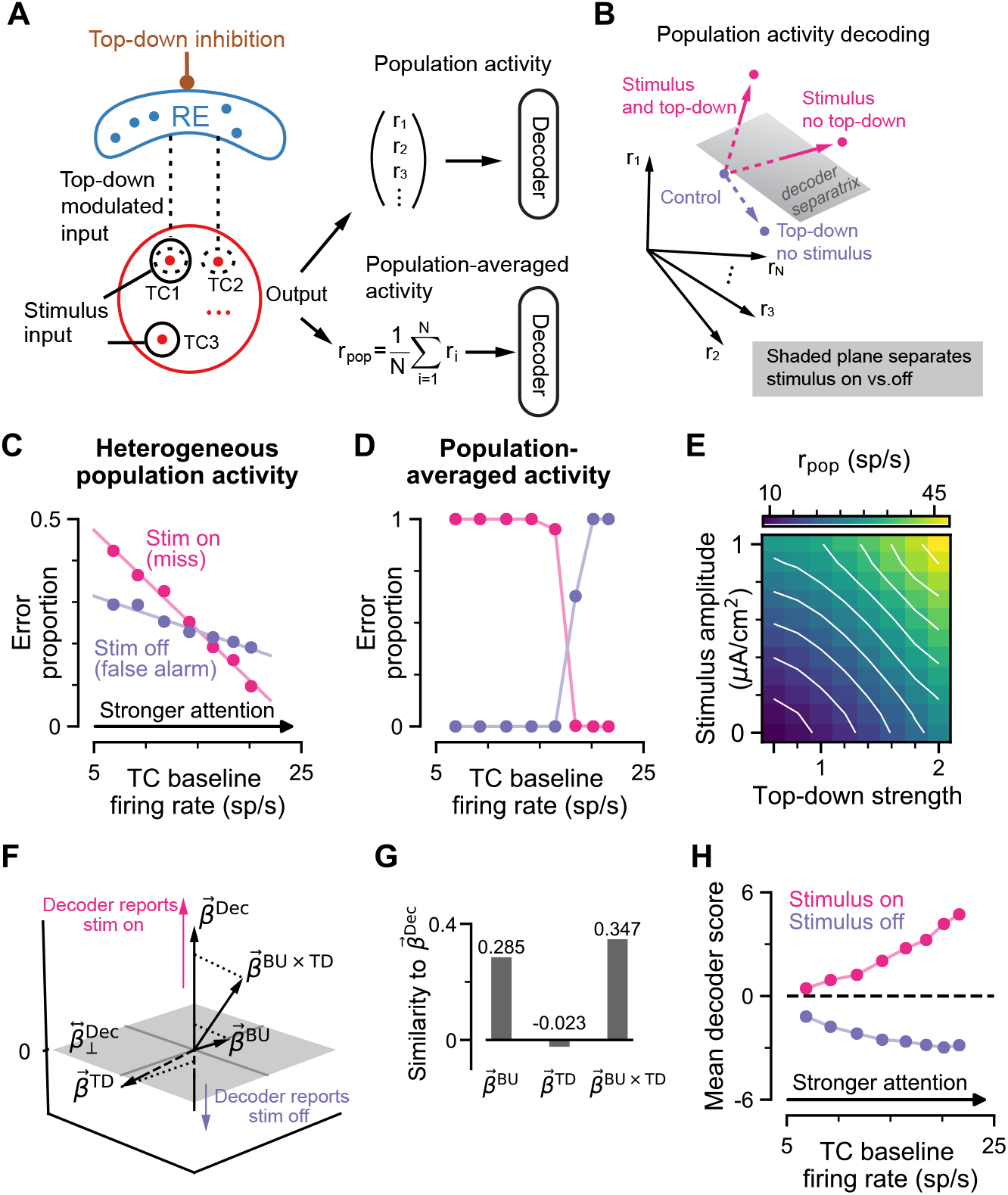
Downstream decoding utilizes heterogeneous population activity patterns for detection. **(A)** Schematic of readout via population decoders. Heterogeneity of circuit properties leads to inter-neuronal variation in net input to TC neurons from bottom-up stimulus and top-down (mediated by RE) pathways. Stimulus on vs. off conditions be decoded from high-dimensional TC population activity patterns (top), or alternatively via the mean activity (*r*_pop_) averaged over the TC population (bottom). Here decoders are implemented as linear support vector machines (SVMs). **(B)** Schematic of population-activity decoding in the detection paradigm. The SVM decoder corresponds to a hyperplane separatrix that maps activity patterns on each side of the hyperplane to “stimulus on” vs. “stimulus off” reports. **(C)** For the heterogeneous population activity decoder, top-down modulation via RE improves detection performance by reducing error rates for both misses and false alarms. Misses vs. false alarms are error types when the stimulus was on vs. off, respectively. TC baseline activity serves as a proxy for top-down modulation strength **(D)** The population-averaged activity decoder exhibits a trade-off between miss and false alarm rates, as top-down strength is varied. **(E)** TC population firing rate (*r*_pop_) as a function of bottom-up stimulus amplitude and top-down strength. Both factors increase *r*_pop_ in a similar manner, explaining poor performance of the population-averaged decoder. **(F)** Geometric relationships among coding and decoding axes in TC activity space. Shown are regression weight vectors for bottom-up stimulus 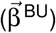, top-down modulation 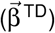, and their interaction 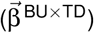, in the 3-dimensional subspace spanned by 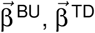, and the SVM decoder weight vector 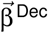 (see **Methods**). **(G)** Projections of 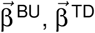 and 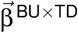 onto 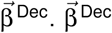 has a positive alignment with 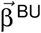 and 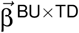, and a weak negative alignment with 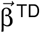. Top-down modulation via RE improves detection performance mainly through gain modulation, evident from the strong positive alignment of 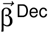 with 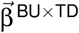. **(H)** The average SVM score (i.e., the projection of the circuit activity onto 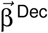), as a function of top-down strength, when the stimulus is present or absent. Top-down modulation shifts scores for both conditions away from the classifier boundary at 0, leading to reduced rates of misses and false alarms (in **C**).

We first varied the top-down strength and characterized its impact on different types of errors in the detection task. From signal detection theory, decoder errors can be decomposed in false alarms (i.e., responding ‘yes’ when the stimulus was absent) and misses (i.e., responding ‘no’ when the stimulus was present) (**Figure S4A**). The decoder based on population-averaged activity exhibited a sharp tradeoff between misses and false alarms as attentional strength varied (**Figure 4D**), in a manner consistent to classic signal detection theory. This tradeoff arises in the population-averaged decoder because stimulus input and top-down modulation both increase population averaged activity (**Figure 4E**). Remarkably, we found that strengthening top-down modulation reduced both types of errors for the decoder based on heterogeneous population activity patterns (**Figure 4C**). Under stronger top-down modulation, TC baseline activity is elevated yet there is decreased probability of downstream misclassification of that activity pattern as stimulus presence. These findings show that the improvement in downstream readout due to modulation of a distributed code across TC neurons, rather than their aggregate activity level.

To gain further insight into the neural coding principles for attentional enhancement of performance, we examined geometric relationships of coding axes in the high-dimensional space of TC activity patterns (**Figures 4F–H, S4**). The weight vector of the SVM decoder defines an axis in activity space, perpendicular to the classifier hyperplane, that assigns a score to each activity pattern by its projection along the decoder axis. In addition, we used regression analysis to define other weight vectors related to three aspects of TC activity: (i) bottom-up, reflecting the stimulus; (ii) top-down, reflecting the influence of the attention signal itself; and (iii) the interaction of bottom-up and top-down, reflecting attentional gain-like modulation of stimulus representations.

As expected, the bottom-up weight vector has a positive cosine similarity with the decoder axis, so that stronger stimuli increase decoder scores (**Figure 4F–H**). Interestingly, the top-down weight vector has a weak negative cosine similarity with decoder axis. In the absence of bottom-up stimulus, stronger top-down modulation thereby yields stronger negative decoder scores, which reduces false alarms. Finally, the bottom-up–top-down interaction weight vector has a strong cosine similarity with the decoder axis. Gain-like modulation by top-down signals thereby increases decoder scores during stimulus presentation, which reduces misses. These functional analyses reveal how top-down modulation can increase the separability of TC activity patterns for stimulus on vs. off conditions to improve behavioral performance. These geometric properties also provide predictions for analysis of empirical TC spiking activity during attentional tasks.

### Attentional Filtering of Distractors to Prevent Interference

In the following, we considered the important attentional function of flexibly regulating stimulus processing across multiple sensory modalities according to task demands (**Figure S3**). In this study, we specified two stimulus modalities, with one as the target for attention, and the other as a distractor. The stimulus for each modality is sent to a separate TC-RE circuit (i.e., signals for two modalities enter separate thalamic nuclei and associated TRN regions). To apply attentional control, RE neurons for the attended modality will be strongly inhibited by top-down input, while RE neurons for the unattended modality will be only weakly inhibited. This allows the signal from the attended modality to be amplified, and that from the unattended modality to be suppressed. Here we controlled the attentional bias by parametrically varying the top-down strength to the distractor modality. Importantly, the stimuli from the two modalities can be congruent, such that both stimuli consistent in their mapping to a response, or they can be incongruent, such that the two stimuli map to opposing responses (Ahrens et al., 2015; Wimmer et al., 2015). We found that in the cross-modal paradigm, the circuit model demonstrates higher discrimination accuracy for congruent trials than for incongruent trials (**Figure S3H-I**). Strong attentional bias dramatically improves performance on incongruent trials, because the distractor stimulus signal is suppressed and filtered out relative to the target signal. This filtering also attenuates the performance benefit from the distractor in congruent trials. Attentional bias results in a performance tradeoff, although congruent trials still have higher sensitivity than incongruent trials and the net effect of attentional bias in improved performance. These findings suggest top-down modulation can implement flexible cross-modal filtering out of irrelevant signals to prevent interference with contextual behavior based on attended signals.

### Sensitivity Analysis of Biophysical Perturbations

How are circuit dynamics and computations shaped by neuronal and synaptic properties? We performed a sensitivity analysis on the spiking circuit model, measuring how strongly a circuit feature is changed by a small perturbation of a given parameter. This analysis characterizes along which directions in parameter space the circuit features are sensitive or robust, how different parameters could compensate for perturbations, which features are sensitive or robust, and how features covary under perturbation. Specifically, we perturbed 6 biophysical parameters (4 synaptic and 2 cellular) (**Figure S5A**), and measured 7 features (3 dynamical and 4 task-computational), generating a feature sensitivity matrix (**Figure S5B**). Averaging across all features, the RE-specific parameters generated greater feature sensitivities than did TC-specific or synaptic parameters (**Figure S5B** top). In further analysis using singular vector decomposition (SVD, see **Methods**), we found that that the circuit’s sensitivity to perturbations is approximately one-dimensional (**Figure S5C**), and TC- and RE-related perturbations have opposite-sign scores, with synaptic perturbations at intermediate scores (**Figure S5D**). These findings revealed that the dominant mode of sensitivity can be characterized as capturing alteration in excitation-inhibition balance.

### Geometrical Relationships of Population Coding in Recordings from Auditory Thalamus

As described in the above sections, our circuit model makes predictions for population coding of bottom-up and top-down signals in sensory thalamus during attentional tasks. Motivated by these model predictions, we trained animals on the cross-modal divided attention task which we had previously described (Wimmer et al., 2015; Nakajima et al., 2019). Briefly, on each trial, an animal was presented with a cue (100 msec of high pass or low pass noise burst) which indicated which one of two target modalities (vision vs. audition) to select. Following a 500-700 msec delay period, a light flash and auditory sweep were simultaneously presented. The light flash location signaled the appropriate port to collect the reward on *attend to vision* trials, whereas an up- or down-sweep indicated whether the right or left port was the reward location on *attend to audition* trials (**Figure 5A**). Following training to criterion (see **Methods**), animals were implanted with multi-site multielectrode arrays targeting the auditory thalamus (ventral medial geniculate body [MGBv]) and primary auditory cortex (A1).

**Figure 5:**
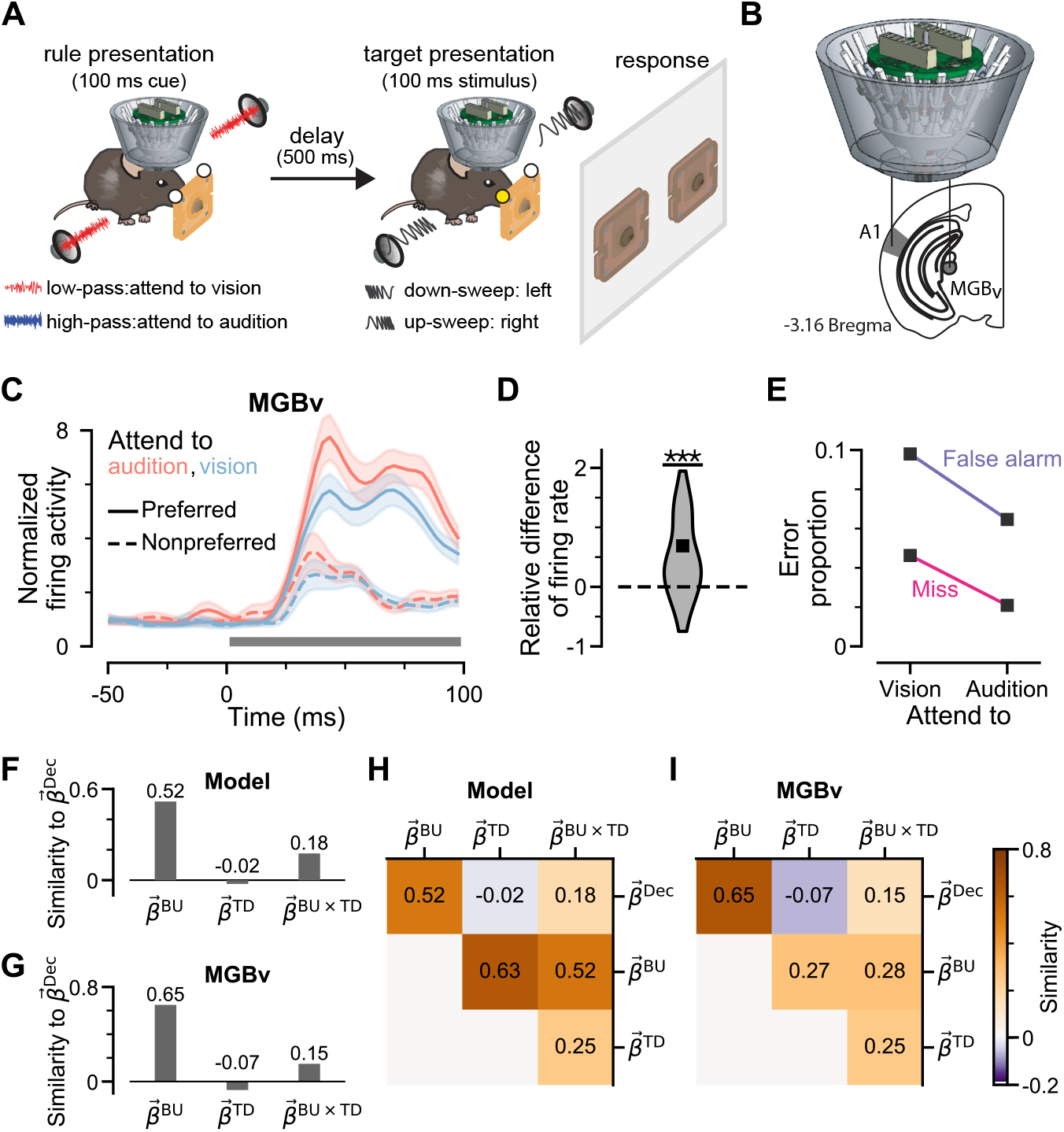
Geometrical coding properties in auditory thalamus (MGBv) recordings. **(A)** Schematic of cross-modal sensory selection task. After self-initiation, freely behaving mice are cued by either high-pass or low-pass noise, which correspond to two rules, “attend to vision” or “attend to audition”, respectively. Following a 500-ms delay, two sensory targets — 100 ms of a visual flash and auditory sweep — are simultaneously presented to indicate the rewarded port location for the cued modality. **(B)** Schematic of multi-electrode drive targeting auditory thalamus (ventral medial geniculate body [MGBv]) and primary auditory cortex (A1). **(C)** Normalized firing rates averaged across all MGBv neurons (n=52 neurons from 2 mice). Top-down attention to the associated modality, audition, increases spiking activity to preferred stimuli more than to nonpreferred ones. The gray bar indicates stimulus presentation. Shaded areas show standard error of mean (s.e.m.). **(D)** Distribution across MGBv neurons of the relative change of evoked activity difference between audition and vision attentional conditions. The relative difference is defined as the firing rate change for “attention to audition” minus that for “attend to vision”, normalized by that for “attend to vision”. The firing rate change was calculated as the response to preferred stimulus minus that to non-preferred. Attention to audition increases difference between preferred and nonpreferred stimuli (Wilcoxon test, p<0.001). **(E)** Error proportion as a function of attended modality for the SVM decoder. Both misses and false alarm errors were reduced when the attention directed to the corresponding sensory modality. **(F,G)** Cosine similarity between the decoder and regression weight vectors obtained from model simulations **(F)** and MGBv recordings **(G)**, which are similar to each other and consistent with results in **Figure 4G**. **(H,I)** The similarity between the upper triangular cosine similarity matrix (which characterizes relative orientations of each vector) from model simulations **(H)** and MGBv recordings **(I)** is 0.77.

In this task, we had previously found that MGBv neurons exhibit a preference toward a specific auditory stimulus (up-versus down-sweep) (Nakajima et al., 2019). In order to relate these empirical MGBv recordings to model predictions in detection paradigms, we grouped the firing activities of preferred stimulus and the firing activities of nonpreferred stimulus for all MGBv cells at the two attentional conditions. The firing patterns of the preferred stimulus were used to mimic stimulus-on trials, whereas those of nonpreferred stimulus were used to mimic stimulus-off trials. As shown in **Figure 5C**, the firing activities of MGBv neurons showed a stronger response during *attend to audition* trials. The response gain, quantified here as the difference between evoked responses to preferred and nonpreferred stimuli, were also larger when the attention was directed toward audition (**Figure 5D**). Higher SVM decoder accuracy indicates that MGBv neurons emitted spikes that were more informative of these targets during the *attend to audition* compared to *attend to vision* (**Figure S6A**). Further analysis revealed that attentional improvement in decoding accuracy was due to reduction of both miss and false alarm error types (**Figures 5E, S6B-C**). The findings are consistent with our model predictions (**Figure 4C**), in that the decoder can distinguish between firing-rate increases due to bottom-up stimuli and those from top-down attentional modulation.

We next calculated population coding axes in the empirical MGBv spiking activity, using the regression modeling framework applied to circuit model, to obtain the bottom-up, top-down, and interaction coding directions as weight vectors in population state space. The geometrical relationships between different weight vectors were found to be consistent between MGBv recordings and our circuit model results (**Figure 5F-I**). Specifically, the bottom-up and interaction weight vectors have positive similarities with the decoder vector, whereas the top-down weight vector has a negative similarity (**Figure 5F-G**). As shown in the model (**Figure 4F-H**), this negative similarity of the top-down vector enables attention to reduce false alarms by decreasing the decoder score when stimulus is absent, while positive similarity of the interaction vector enables attention to reduce misses by increasing the decoder score when stimulus is present.

### Coding Properties in Primary Auditory Cortex

In addition to our analysis of recordings from MGBv, we examined the coding properties of A1, the sensory cortical area directly downstream of MGBv. Comparative analysis of A1 with MGBv enabled us to investigate two main issues. First, because MGBv receives corticothalamic feedback projections from A1, it is an open question whether attentional gain modulation of thalamic activity is computed locally within thalamus, as proposed by our circuit model mechanisms, or instead whether the gain modulation could be inherited from activity in A1 (**Figure 5A**). Second, our SVM analyses of model and empirical MGBv activity suggest how downstream decoding from thalamus could separate bottom-up from top-down signals, and signatures of such a readout mechanism can be tested in A1 activity.

As with MGBv, A1 neurons showed to have higher firing rates when attending to audition (**Figure 6B**). In order to investigate differences between MGBv and A1 neurons in attentional gain modulation, we calculated the gain modulation factor as the interaction term divided by the bottom up term from the regression model, which captured the attentional effect on the bottom-up strength. We characterized the time courses of gain modulation in MGBv and A1, and found earlier gain modulation in MGBv than in A1 (**Figure 6C–D**). These timing results appear inconsistent with gain modulation in MGBv being inherited from A1, and suggest that thalamic gain modulation results from the local TC-RE circuit computations rather than cortical feedback, at least for this particular task (**Figure 6A**).

**Figure 6:**
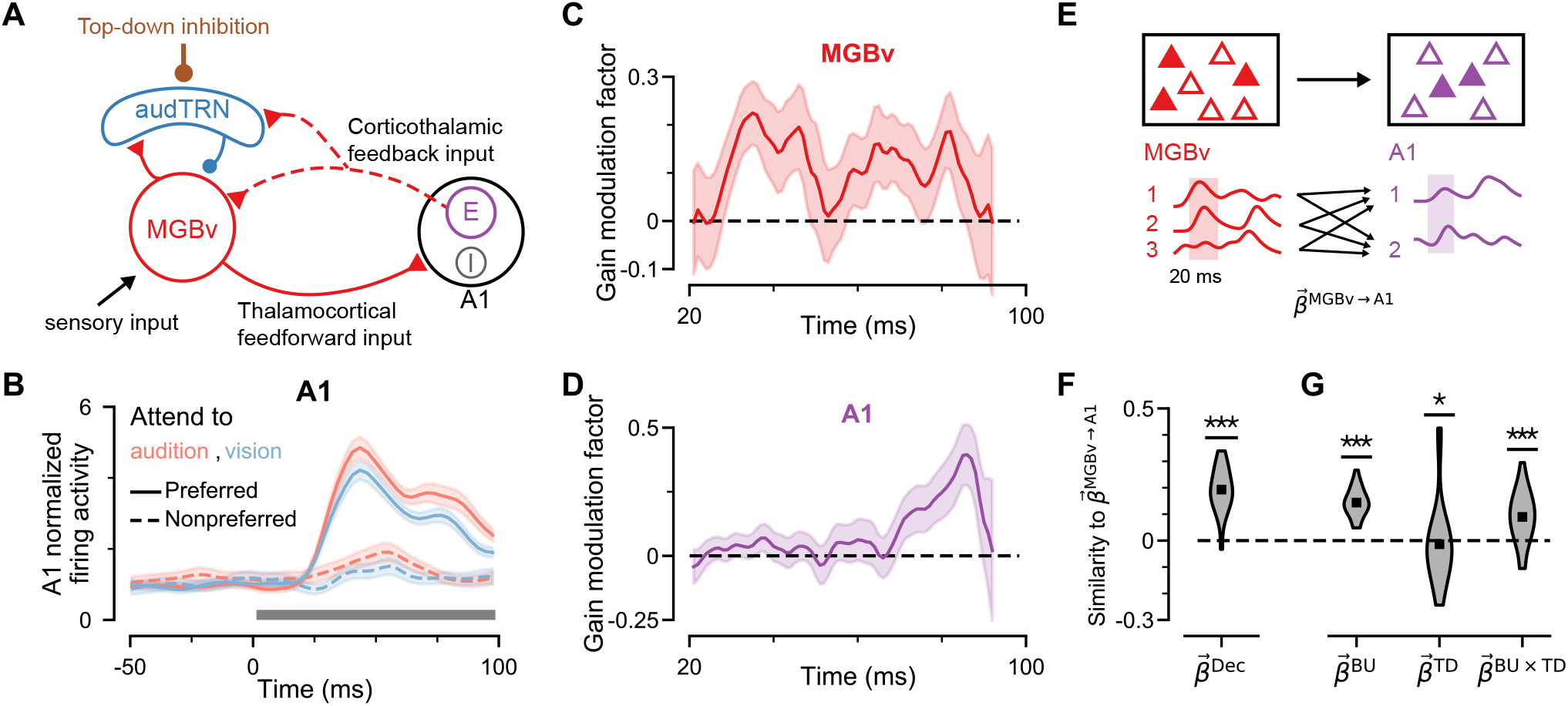
Coding properties of primary auditory cortex compared to thalamus. **(A)** Schematic of the auditory thalamo-cortical loop. Gain modulation of thalamic activity could be generated within the thalamic circuit, or potentially reflect gain modulation generated within downstream sensory cortex which is transmitted via cortico-thalamic feedback connections (dashed line). **(B)** Normalized firing activity averaged across all A1 neurons (n=25 neurons from 2 mice). The horizontal bar indicates stimulus presentation. **(C,D)** Average gain modulation factor of MGBv and A1 neurons over time. Significant increase of the A1 gain modulation factor was later than the that of MGBv neurons, indicating that gain modulation in MGBv is not primarily inherited from A1. The shaded area represents s.e.m. **(E)** Schematic of mapping from source MGBv activity to target A1 activity. Filled triangles indicate active neurons. Mapping weights 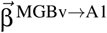 were obtained by a linear regression approach (see **Methods**). **(F,G)** Cosine similarity of the mapping weights from MGBv to A1 neurons to the decoder axis and three coding axes (from **Figure 5G**). Mapping weights have a positive similarity with the decoder axis **(F)**. Meanwhile, mapping weights have a positive, negative, and positive similarity with bottom-up, top-down, and interaction axis, respectively (Mann-Whitney U test, decoder: p<0.001; bottom-up: p<0.001; top-down: p=0.025; interaction: p<0.001) **(G)**.

We next further analyzed the peri-stimulus-time histograms (PSTHs) of A1 and MGBv during stimulus presentation. We examined how neuronal activity in A1 was related to that in MGBv, using regression-based approaches (Semedo et al., 2019; 2020; Keeley et al., 2020) (**Figure 6E**, see **Methods**). Specifically, we calculated the mapping weights to reconstruct each A1 neuron’s PSTH from those of MGBv neurons under different conditions. Each A1 neuron’s weight vector can be thought of as effective readout weights from MGBv for that A1 neuron. We found that weight vector of each A1 neuron and the SVM weight vector from MGBv firing activity had a positive cosine similarity (**Figure 6F**). Moreover, the weight vector of each A1 neuron has significant positive, negative and positive cosine similarity with the bottom-up, top-down and interaction vector, respectively (**Figure 6G**). The signs of these cosine similarities are preserved when incorporating time lags, or using Poisson generalized linear model (**Figure S6D–G**). The relationships are consistent with those among the SVM weights and the three coding axes in MGBv firing activities and predicted by our model (**Figure 5F–I**). Therefore, these results support the plausibility of such a decoding axis from MGBv to A1, through which neurons can distinguish bottom-up from top-down signals.

### Effects of Recurrent Excitation on Population Coding

A distinctive characteristic of thalamic microcircuitry is the lack of recurrent excitatory synaptic connections among TC neurons (Jones, 2012), in contrast to highly recurrent cortical circuits (Ozeki et al., 2009; Sanzeni et al., 2020). Here we extended our models to investigate how recurrent excitation affects population coding of bottom-up and top-down signals, and how this impacts downstream decoding (**Figure 7A**). We considered a thalamic circuit consisting of large number of heterogeneous rate units matched to the spiking model (see **Methods**). We parametrically varied the strength of recurrent excitation among TC neurons, and found that the similarity between bottom-up or top-down coding axes (BU-TD similarity) increases when with stronger recurrent excitatory coupling *J*_EE_ (**Figure 7B**). Increasing recurrent excitatory coupling strength *J*_EE_ can thereby reduce separability of BU and TD signals in the TC population. Empirical analysis of MGBv and A1 recordings revealed results consistent with this model prediction: A1 neurons have a higher BU-TD similarity than MGBv neurons (**Figure 7C**). In addition, we applied an SVM decoder to the model and found higher decoding accuracy when recurrent excitation is weaker (**Figure 7D**). These results suggest functional advantages for thalamus, lacking recurrent excitation, as a locus for top-down attentional control. They further highlight a unique aspect of a modeling approach when, to our knowledge, the ability to experimentally introduce recurrent excitation to the intact biological thalamus is not possible.

**Figure 7:**
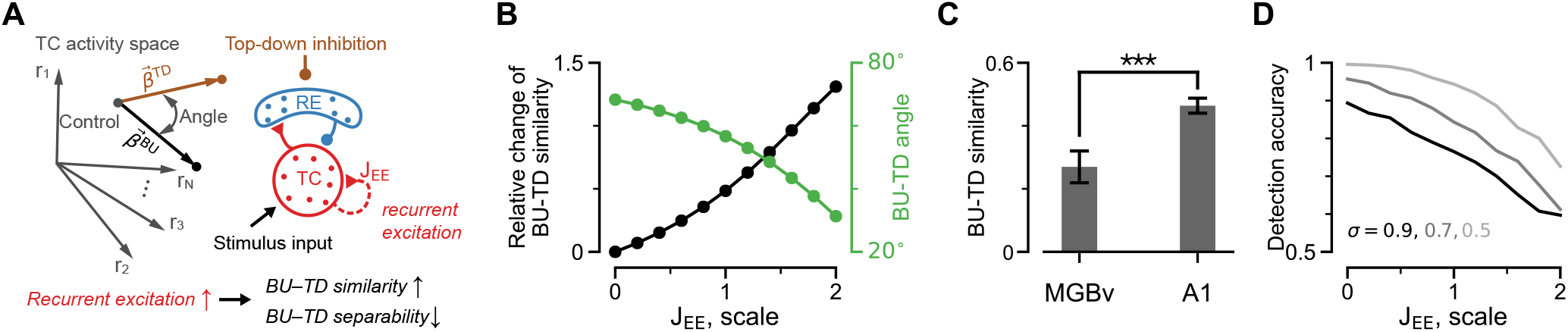
Recurrent excitation degrades separability of population coding of bottom-up and top-down signals. **(A)** Schematic of TC activity space. Recurrent excitation among TC neurons (*J*_EE_) affects the similarity, and separability, of bottom-up (BU) and top-down (TD) signals in TC population activity. Lower separability of BU and TD representations is reflected in higher similarity of their coding axes with a smaller angle between them. **(B)** Relative change of BU-TD similarity, and the angle between BU and TD coding axes, as a function of recurrent coupling strength *J*_EE_ among TC neurons. Increasing recurrent excitation increased the similarity, and reduced the angle, between BU and TD coding axes. **(C)** Empirical characterization of BU-TD similarity in MGBv and A1 recordings (n=52 MGBv neurons and n=25 A1 neurons, from 2 mice). Error bars denote standard error calculated by jackknife procedure. BU-TD similarity was higher in A1 than in MGBv (t-test, p<0.001). **(D)** Detection accuracy in the model, from an SVM decoder, as a function of recurrent coupling strength *J*_EE_. Detection accuracy decreases when increasing recurrent excitation strength (darker lines denote stronger background noise, with unit μA/cm^2^).

## Discussion

In this study, we developed a biophysically grounded, yet parsimonious, model of a thalamic circuit, which is constrained by experimental findings to operate in the dynamical regime observed in alert behaving animals. The circuit model captures a number of key experimentally observed phenomena related to the thalamus and attention, including gain-like modulation of responses controlled by top-down control signals. Attentional modulation resulted in improved performance for downstream readout in detection, discrimination, and cross-modal task paradigms, with heterogeneity of thalamic response being critical for downstream decoding. Dynamical systems analysis reveals mechanisms by which the inhibitory TRN acts a preferentially potent site for attentional control. Moreover, analysis of neuronal recordings from sensory thalamus and cortex of task-performing animals revealed similar coding axes, and also suggests that the gain modulation in thalamus likely results from local intra-thalamic processing with little contribution from cortical feedback. Finally, our modeling shows how lack of recurrent excitation within thalamus can be beneficial for separability of top-down and bottom-up signals, which was supported by measured separability in neuronal recordings.

Gain modulation of neuronal responses can be found across cortical and subcortical brain areas and is associated with various cognitive functions including attention (Salinas and Thier, 2000; Ferguson and Cardin, 2020). In different circuits, gain control may be mediated by mechanisms such as synaptic noise (Ly and Doiron, 2009), active dentrites (Mehaffey, 2005) and diversity of interneuron populations (Litwin-Kumar et al., 2016). That the thalamic gain is more potently regulated by top-down modulation of RE neurons in the TRN, compared to direct modulation of TC neurons, is a key finding from our model. Interestingly, disinhibitory motifs are also prominent in neocortical microcircuits, and this computational principle may support contextual modulation of cortical state and response gain (Kepecs and Fishell, 2014; Fu et al., 2014; Litwin-Kumar et al., 2016; Yang et al., 2016).

Heterogeneity of thalamic activity patterns plays a critical role for attentional modulation to enhance downstream readout of stimulus information. Because both bottom-up and top-down inputs increase TC mean firing rates, readout of aggregate activity cannot disambiguate stimulus from attention signals. In the heterogeneous spiking circuit model, TC neurons are differentially responsive to bottom-up stimulus, top-down modulation, and their gain-like interaction, which allows downstream readout to harness attentional gain modulation to improve performance in detection and discrimination paradigms. These features could not be captured in homogeneous or mean-field models of thalamus (Roberts and Robinson, 2012; Jaramillo et al., 2019). Importantly, we found similar geometrical relationships of coding axes in the model and in empirical MGBv neurons. However, note that, the tasks from which we obtained the MGBv recordings did not vary stimulus difficult near psychophysical thresholds as in typical detection paradigms (Nakajima et al., 2019). To more fully test the model predictions, future experiments should record from thalamus during cross-modal detection tasks that manipulation stimulus magnitudes and attentional states. Interestingly, the model shows degraded separability of multiplexed bottom-up and top-down signals, with impaired downstream decoding, if strong recurrent excitation is included, which suggests an advantage of thalamus, compared to cortex, as an effective target for top-down control.

The circuit model developed here is parsimonious, including key biophysical elements, and provides opportunities for extension to address additional neurobiological questions in future computational studies. For instance, models can be extended to incorporate the rich diversity of short-term synaptic plasticity dynamics at their inputs and local synaptic connections (Sherman, 2012; Usrey et al., 1998; Crandall et al., 2015). Furthermore, the TRN contains subcircuits involving distinct inhibitory cell types with different neuronal and synaptic properties (Ahrens et al., 2015; Clemente-Perez et al., 2017; Li et al., 2020; Martinez-Garcia et al., 2020). The model can also be extended to study regional specialization of TC-RE microcircuitry, which exhibits a sensory-to-higher-order hierarchical organization (Phillips et al., 2019; Wei et al., 2011; Li et al., 2020; Martinez-Garcia et al., 2020).

Our sensitivity analysis characterized excitation-inhibition balance as a key effective parameter regulating thalamic dynamics and computation, with preferential sensitivity to perturbations involving RE neurons. The TRN has emerged as a key site of vulnerability associated with neurodevelopmental disorders and as a target for therapeutics (Krol et al., 2018; Huguenard and McCormick, 2007; Fagerberg et al., 2014; Ritter-Makinson et al., 2019; Ahrens et al., 2015; Wells et al., 2016; Murray and Anticevic, 2017; Richard et al., 2017). Incorporation of additional biophysical processes in the model will likely be important for simulating effects of disease and pharmacology, to link cellular and synaptic perturbations to deficits in perceptual and cognitive functions.

Finally, future studies should incorporate biophysically grounded models of thalamus into large-scale brain models of cognitive functions such as attentional control and decision making. The present circuit model can be embedded in networks with long-range interactions involving thalamic, cortical and basal ganglia circuits, to gain further insight into the distributed computational mechanisms of attentional function (Saalmann and Kastner, 2011; Sherman, 2016; Halassa and Kastner, 2017; Nakajima and Halassa, 2017; Rikhye et al., 2018; Schmitt et al., 2017; Krauzlis et al., 2018). Our present study thereby provides a foundation for future computational modeling of cognitive functions involving the distributed interactions of subcortical and cortical circuits

## Acknowledgements

This research was supported by NIH grants R01MH112746 (JDM), R01MH107680 and R01NS098505 (MMH); NSERC (NHL); and the Swartz Foundation (QLG). We thank members of the Murray and Halassa labs for helpful discussions and feedback on the manuscript.

## Author Contributions

Conceptualization: QLG, NHL, RDW, MMH, JDM; Methodology: QLG, NHL, JDM; Software: QLG, NHL; Data Collection: RDW; Formal Analysis: QLG, NHL; Writing – Original Draft: QLG, NHL, JDM; Writing – Review & Editing: QLG, NHL, RDW, MMH, JDM; Visualization: QLG; Supervision: JDM; Project Administration: JDM; Funding Acquisition: JDM.

## Methods

### Spiking Circuit Model

The thalamic circuit model consists of *N*_TC_ = 1000 excitatory thalamocortical (TC) neurons and *N*_RE_ = 1000 inhibitory reticular (RE) neurons. TC and RE neurons are interconnected through sparse and random synaptic projections, without projections between neurons of the same cell type, with probability of connection *p*_TC→RE_ = *p*_RE→TC_ = 0.01 and *p*_TC→TC_ = *p*_RE→RE_ = 0 (Jones, 2012; Hou et al., 2016). Both TC and RE neurons receive external excitatory inputs, and RE neurons additionally receives external inhibitory inputs. All excitatory synaptic connections are mediated by α-amino-3-hydroxy-5-methyl-4-isoxazolepropionic acid receptor (AMPAR) conductances, and all inhibitory synaptic connections are mediated by γ-aminobutyric acid type-A receptors (GABA_A_R) conductances, following:

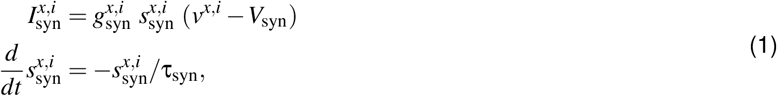

where *x* refers to the target neuron group (TC or RE), *i* refers to the neuron index within the group, and *syn* refers to the synapse type (AMPA or GABA_A_). *V*_AMPA_ = 0 mV, *V*_GABA_ = −80 mV, τ_AMPA_ = 2.5 ms, and τ_GABA_ = 10 ms are the reversal potentials and synaptic timescales of AMPAR and GABAR.

To produce distributions of firing rate and burst rate comparable to experimental observations, the synaptic conductances of external inputs 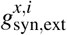 are set to be heterogeneous, with conductance strengths following a log-normal distribution with mean 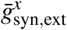 and standard deviation 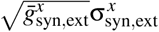. The extent of heterogeneity of the external inputs and the neuronal firing rates in the circuit is thereby shaped by 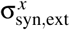. The parameters for external inputs are 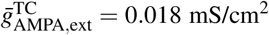, 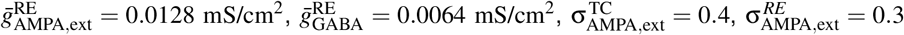 and 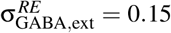. In **Figure S1**, 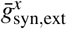 is instead varied so that mean firing rate is fixed around 11 Hz for TC neurons and 16 Hz for RE neurons. The synaptic conductances for connections between TC and RE neurons are constant, with 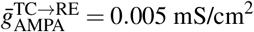 and 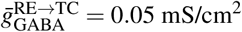.

### TC Neuron Model

The TC neuron is modeled by a single-compartment Hodgkin-Huxley model, with a minimal set of active conductances sufficient o endow the neuron with characteristic the tonic and burst spiking dynamics. The model is similar to prior models of TC neurons (Destexhe et al., 1993b;a), and specific TC channel conductance equations used here were adapted from prior TC neuron modeling (Destexhe et al., 1993a; Golomb et al., 1996). In all equations below describing neuronal currents, membrane potential has the unit of millivolt (mV), and temperature factors *T*_adj_ were set with temperature *T* = 36°C.

The membrane potential of the TC neuron is governed by the following dynamics:

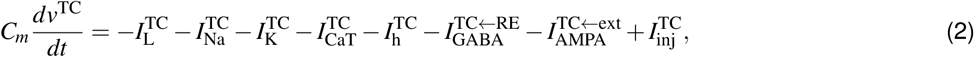

where 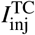 is any injected current input, and 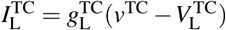 is the leak current with 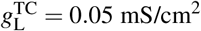 and 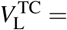 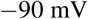. The Na^+^ current is given by :

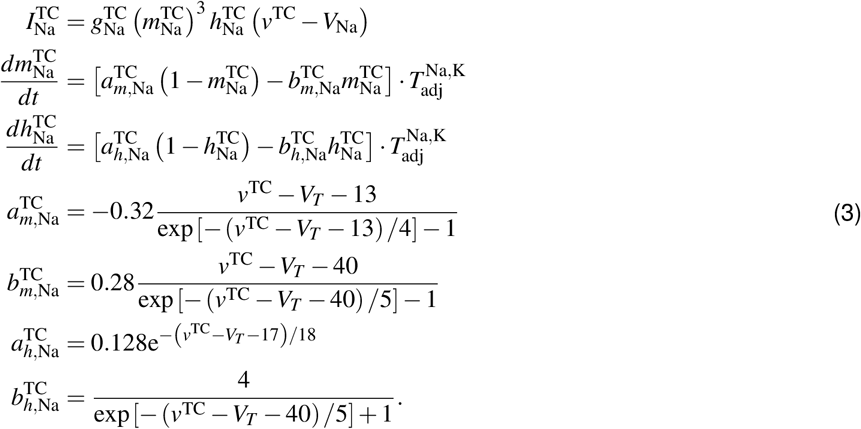

where 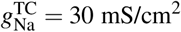, *V_Na_* = 50 mV, and the spike adjusting threshold *V_T_* = −55 mV. The temperature factor is given by 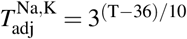.

The K^+^ current dynamics are given by:

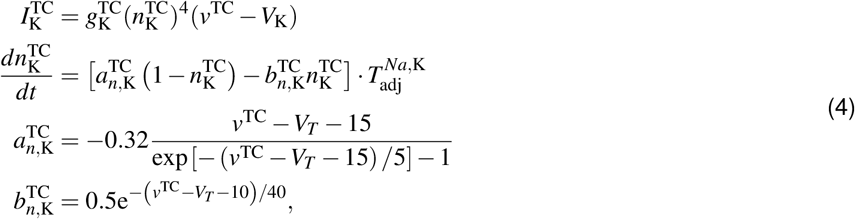

where 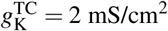, and *V*_K_ = −95 mV.

The low-threshold Ca^2+^ current dynamics are given by:

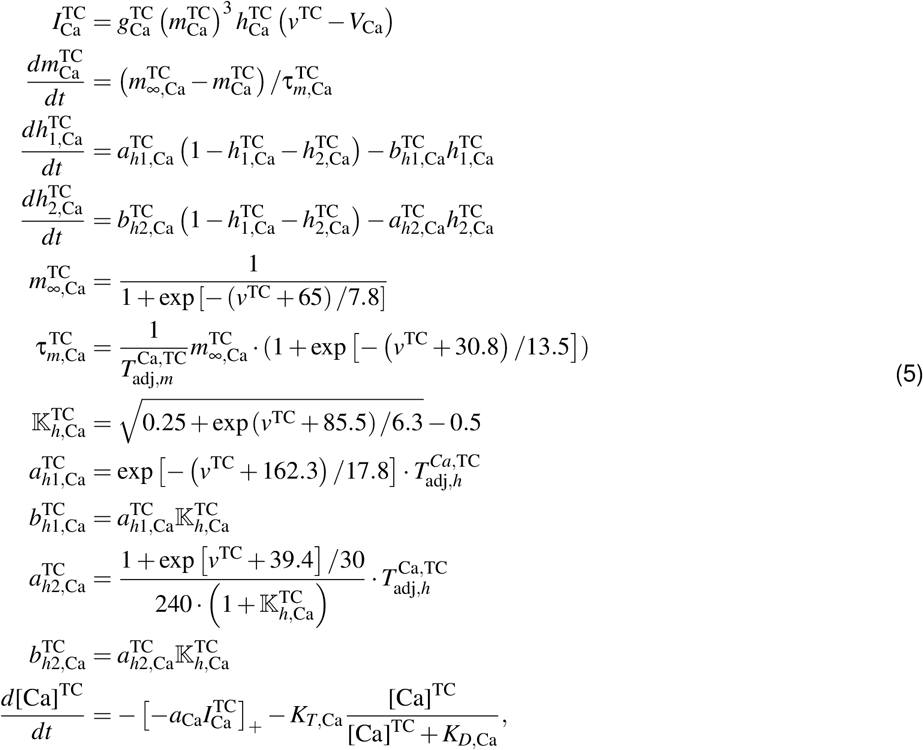

where 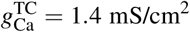, and *V*_Ca_ = 120 mV. The temperature factors 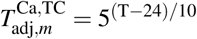 and 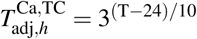. The gating kinetics was adjusted to 36 °C. Parameters for the Ca^2+^ kinetics, *a*_Ca_ = 1/(2 × 96489 Cmol^−1^ × 1μm), *K_T_*_,Ca_ = 0.1μM/ms, and *K_D_*_,Ca_ = 0.1μM.

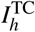 is modeled as a single-variable kinetic scheme with the dependence of its activation on voltage and intracellular Ca^2+^. The kinetic equations are as following:

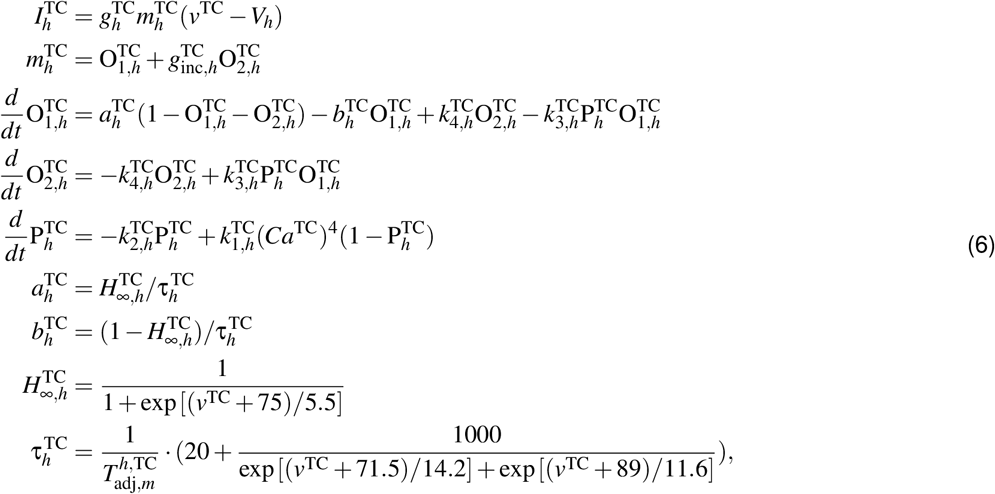

where 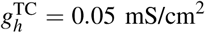, *V_h_* = −43 mV, and 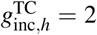. According to the channel properties, the reaction rates are 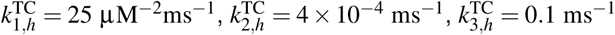, and 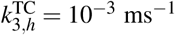.

### RE Neuron Model

The RE neuron is also modeled by a single-compartment Hodgkin-Huxley model, with the specific channel conductance equations adapted from prior RE neuron modeling (Destexhe et al., 1994).

The membrane potential of the RE neuron is governed by the following dynamics:

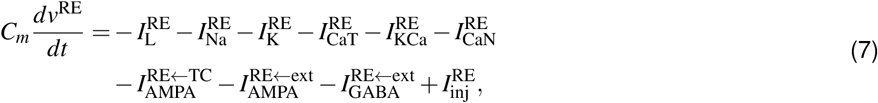

where 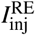 is any injected current input, and the leak current is 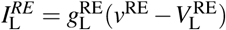 with 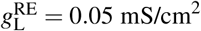 and 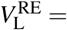 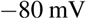. Here 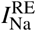 and 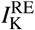 have the form as Eq. (3) and Eq. (4) respectively, with 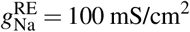 and 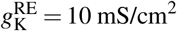.

The low-threshold Ca^2+^ current 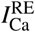 has the following form:

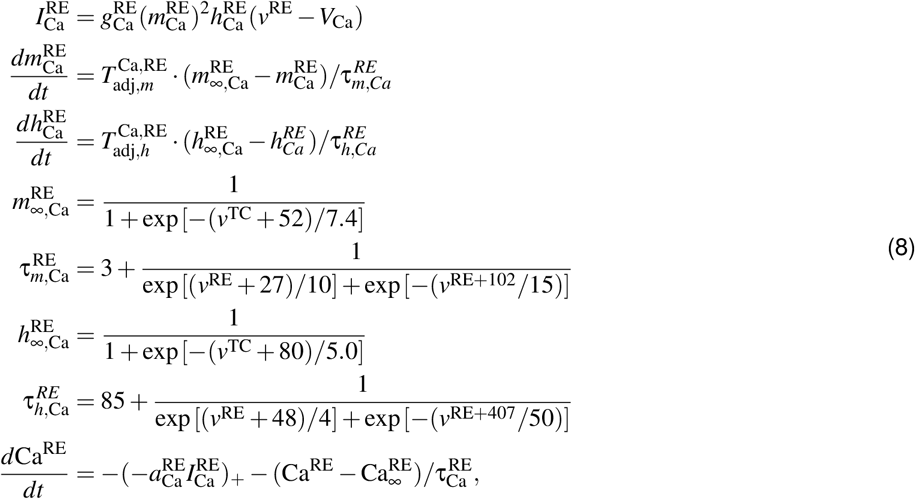

where 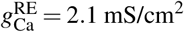. The temperature adjusting terms 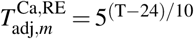 and 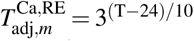. The gating kinetics was adjusted to 36 °C. Parameters for the Calcium kinetics, 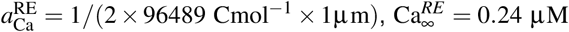, and 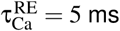.

The Ca^2+^-dependent K^+^ current is described as:

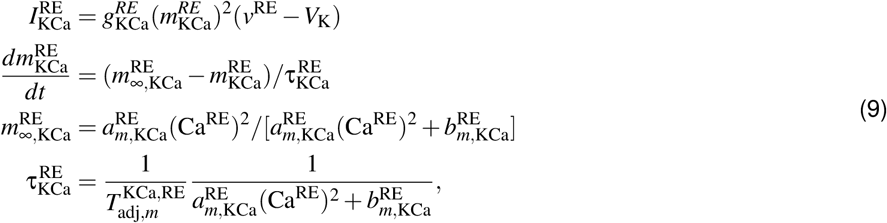

where 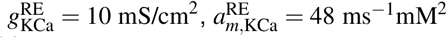. and 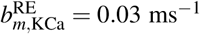. The temperature adjusting term is 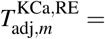 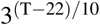.

The Ca^2+^-dependent nonspecific cation current follows the equation:

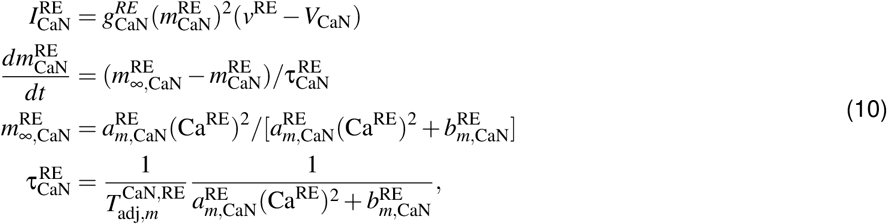

where 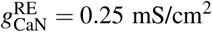, *V*_CaN_ = −20 mV, 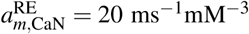, and 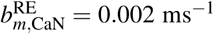, and the temperature adjusting term 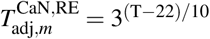.

### Top-Down and Stimulus Inputs

Top-down inputs are stimulated as Poisson-distributed spike trains, of a given rate, driving either GABA_A_ or AMPA synapses on either RE or TC neurons. The external spike trains are generated by a Poisson process with rate 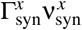 for syn = AMPA, GABA and *x* = TC, RE. Here, 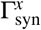 is the factor denoting the strength of the top-down attentional modulation, with 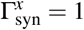 during the baseline state. We set 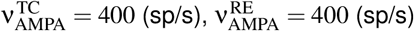 and 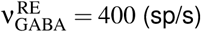 to obtain the the mean firing rate is around 11 (sp/s) for TC neurons and 15 (sp/s) for RE neurons (**Figure S1B-C**).

For parsimony, stimulus inputs are generally simulated as direct current inputs to TC neurons. The probability of a TC neuron receiving a stimulus input is random with probability *p* = 0.5 to reflect stimulus specificity on sensory inputs to TC neurons. In most figures, stimulus inputs are unstructured with equal strength to all recipient TC neurons. However, in **Figure S3** and corresponding parts of **Figure S5** (*S*^dis^,*S*^cong^,*S*^inco^), stimuli inputs are structured across neuron index to reflect tuned sensory inputs in discrimination tasks, with a Gaussian wave of width σ = 30 neurons around the mean (and extending past neuron 1 to neuron 1000 and vice versa). The two inputs for discrimination have a difference in mean location by 30 neurons.

### Measures of Circuit Dynamics

In **Figures S1**, and **S5**, we characterized aspects of circuit spiking dynamics and their dependence on model parameters. A burst (**Figure S1**) of a neuron is defined as multiple (at least two) spikes with an interspike-interval (ISI) of less than 20 ms, preceded by a long period of 100 ms with no spiking. This definition of bursting is similar to prior definitions used in electrophysiological studies of thalamus (Halassa et al., 2011; Wells et al., 2016). Spindle probability is defined as the probability the spindle power reaches a threshold (**Figure S5**). The spindle power is defined as the power within 9 to 15 Hz band-pass filtered from the power spectrum of population spiking activity.

The awake state regime can be defined by two curves in the parameter space of **Figure S1**. The orange line defines the upper bound before which the spontaneous activity becomes synchronous (**Figure S1G**), and the purple line defines the lower bound for RE activation to generate an evoked spindle-like response (**Figure S1H**). The two lines are defined as the equipotential lines where the maximum power spectral density in either case reach a certain threshold. The threshold is chosen by considering the first column in (**Figure S1H**), where TC neurons do not project to RE neurons. Since any oscillations in the current regime of the circuit model come from reverberation between TC and RE neurons, the circuits in the first column in (**Figure S1H**) are in the asynchronous regime. The threshold is defined as the mean plus twice the standard deviation of maximum power spectral density across that column.

### Reduced Two-Variable Mean-Field Model

For the dynamical systems analysis of **Figure 3**, we reduced the thalamic circuit to a two-variable mean-field description, with one dynamical variable representing the activity of each of the excitatory and inhibitory populations (Wong and Wang, 2006). The synaptic gating variables of the TC and RE populations (*s*^α^, α = TC, RE) follow:

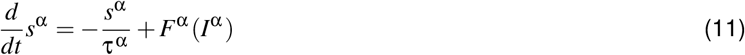

where τ^TC^ = 2.5 ms is the time constant of fast AMPA synapses, and τ^RE^ = 10 ms is the time constant of GABA_A_ synapses. *F*^α^ (·) describes the frequency-current (F-I) curve relation used in prior studies (Abbott and Chance, 2005; Wong and Wang, 2006; Murray et al., 2017):

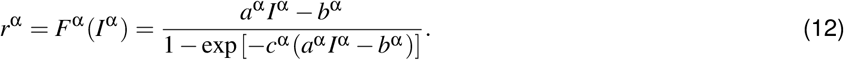

We obtained parameters for the F-I curves by fitting Equation 12 to simulated F-I curves of our Hogkin-Huxley neuron models as shown in **Figure 3B**. Neuronal parameters used with *a*^TC^ = 40.81 (sp/s) · (μ*A*/cm^2^)^−1^, *b*^TC^ = 34.54 (sp/s), and *c*^TC^ = 0.107 s for TC population, as well as *a*^RE^ = 25.97 (sp/s) · (μ*A*/cm^2^)^−1^, *b*^RE^ = −3.91 (sp/s), and *c*^RE^ = 0.222 s for RE population.

The input currents to the TC and RE populations are given by:

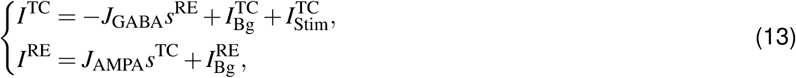

where 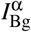 is the background input, and 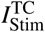 is the stimulus input, *J*_AMPA_ is the coupling strength from TC to RE, and *J*_GABA_ is the coupling strength from RE to TC. Here, we chose the coupling strengths in the mean-field model to be consistent with the spiking circuit model. In the spiking circuit, each TC (RE) neuron receives 10 RE (TC) connections on average with synaptic conductance 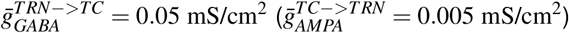. Taking into account the reversal potentials of the AMPA and GABA synapses to estimate approximate ranges of currents, we set *J*_AMPA_ = 4.0 (μ*A*/cm^2^) and *J*_GABA_ = 4.5 (μ*A*/cm^2^). Finally, the background inputs 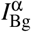 were set so that the TC firing rate is 10 (sp/s) and RE firing rate is 15 (sp/s), in line with the spiking circuit model, with values 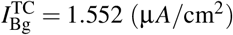 and 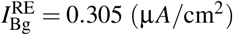.

From eq. (13), we can derive

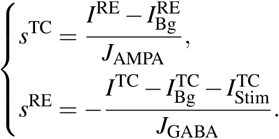

Therefore, eq. (11) can be expressed as

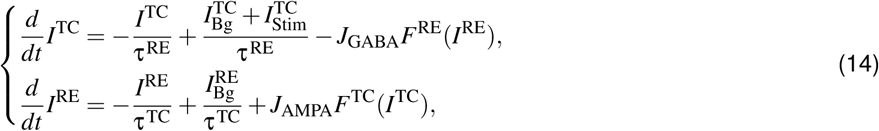

with steady-state solution

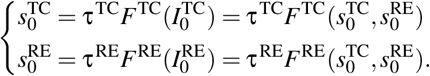

To calculate the response gain of the circuit, we consider the impact of small increment in stimulus input δ *I* to 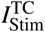. The response gain of the circuit can be defined as the local slope *k*^TC^ for the change in TC firing rate (δ *r*^TC^) resulting from the stimulus increment: δ *r*^TC^ ≈ *k*^TC^δ *I*.

As a consequence of the stimulus increment δ *I*, 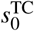 and 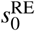 change with increments δ *s*^TC^ and δ *s*^RE^, respectively. Then we can derive the increments of the total current inputs as δ *I*^TC^ = −*J*_GABA_δ *s*^RE^ + δ *I* and δ *I*^RE^ = *J*_AMPA_δ *s*^TC^, and the first-order linearized approximation of the F-I curve as

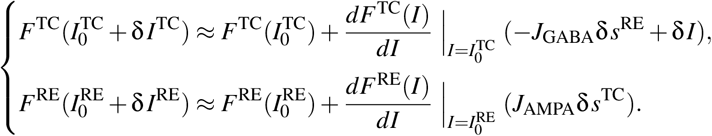

The resulting dynamics of the linearized systems follow

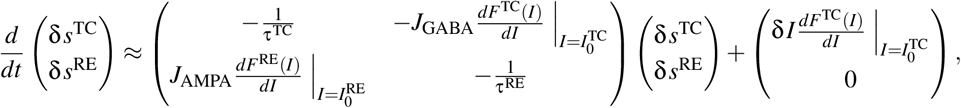

from which we can derive the steady-state increments 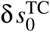 and 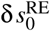 as

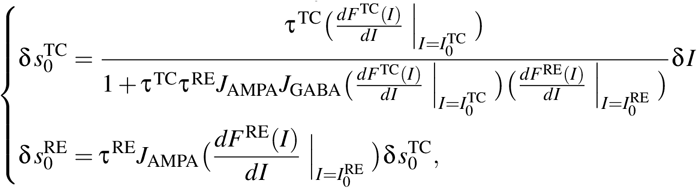

and the increment of TC firing rate as

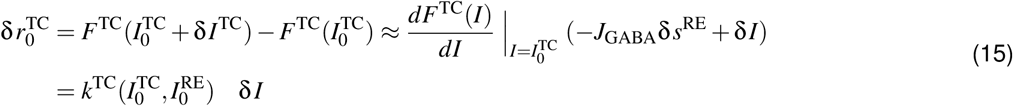

where the circuit response gain 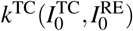 is given by

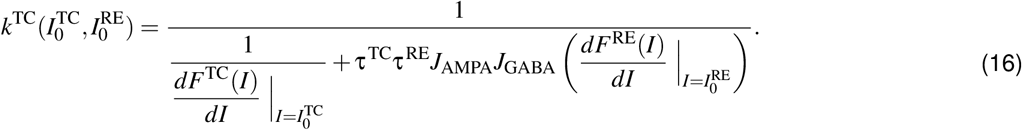

Equation 16 shows that The response gain of the circuit, 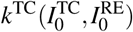, depends on the first derivatives of the F-I curve of both TC and RE neurons. Note that the response gain increases when 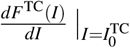 increase or 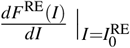 decreases. In addition, the nonlinearity of the F-I curve (usually concave upward) makes 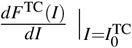 and 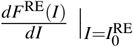 increase when the total current input I increases. Then, in the case of RE modulation, top-down modulation decreases the total input of the RE neurons for the attended modality and suppresses their firing rate, thus increasing the TC firing rate via disinhibition. In this case, 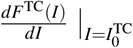 increase and 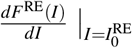 decreases which both increase the slope of the gain curve. In contrast, in the case of TC modulation, top-down modulation increases the total input of TC neurons and amplifies its firing rate, which in turn increases the firing rate of the RE neurons (according to Eq. (13)). As such, both 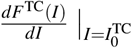 and 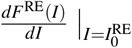 increase, which have opposing and partially canceling-out effects on the response gain. For interpretability and simplicity, **Figure 3** are shown in terms of the firing rates of TC and RE populations, though an equivalent description can be expressed using the synaptic gating variables in the system.

In **Figure 7**, we considered a thalamic circuit of 1000 TC and 1000 RE rate neurons described by **Equation** (11), to characterize the effects of recurrent excitation on population coding of bottom-up and top-down signals. We used the same connectivity matrices as those in spiking networks. We then approximated the target heterogeneous baseline firing rates of TC (RE) neurons by a lognormal distribution with mean 10 sp/s (15 sp/s) and standard deviation 6 sp/s (i.e., similar to that in the spiking network from **Figure S1B**). Then, the recurrent input (inhibition from RE to TC and excitation from TC to RE) was calculated with the previously derived coupling strength, and the background input into each TC or RE neuron was therefore determined with use of the f-I equation. In addition, when investigating the effect of recurrent excitation, we choose τ^TC^ = 10 ms in order to make the system have a stable fixed point and avoid the oscillatory dynamics. Also, we ensure the similar firing activities at the equilibrium state by fixing the product values of τ^TC^*J*_EE_ and τ^TC^*J*_AMPA_.

To be more specific, the total input into the *i*th TC or RE neuron is

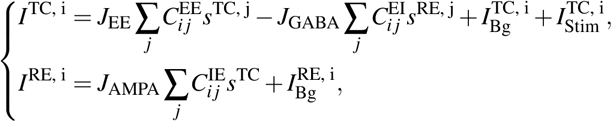

where, *C*^αγ^ for α, γ = E, I are connectivity matrices, which are randomly generated with given probabilities. First we assume 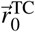 at the equilibrium. There are two kinds of input, bottom-up stimulus 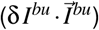 into E population, and top-down inhibition 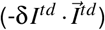 into I population, where δ *I^bu^* and δ *I^td^* are the magnitude of the inputs. 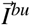 and 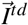 are the normalized vectors with elements decided according to the stimulus and top-down inputs used in the spiking network model in **Figure 7**. However, 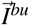 and 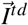 can almost be any normalized vectors with non-negative elements. Supposing with the two inputs, one can obtain the corresponding TC firing rate patterns 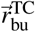 and 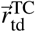. Then their effects can be approximated by

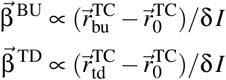

Then with small δ *I*, we can have

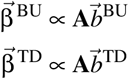

where **A** is matrix dependent with *J*_EE_, i.e. 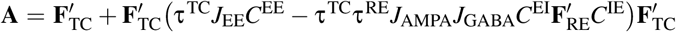. The diagonal matrix 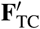 has 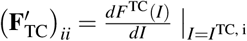. Same as the The diagonal matrix 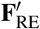. The vector 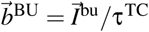, and 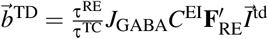. Then with a small perturbation δ_EE_ on *J*_EE_, and assuming small change in the derivative of f-I curve due to this perturbation, one can find

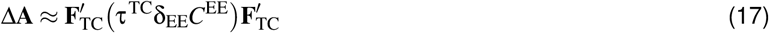

Since *C*^EE^ is a random matrix, 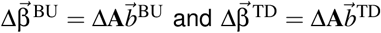 represent two non-negative vectors that are nearly parallel to each other. That is, increase *J*_EE_ can reduce the angle between the two vectors, while amplifying their similarity. Intuitively, increasing recurrent excitation can increase a same amount of input into all the neurons, which reduces the heterogeneity between each neuron.

### Support Vector Machine (SVM) Decoder

To study how stimulus information can be read out from the spiking circuit model, and how its performance can be regulated by top-down inputs, we trained linear support vector machine (SVM) classifiers to perform detection and discrimination. All analysis is conducted using the python package scikit-learn (function linearSVM).

For each task, the firing rate of each TC neuron was calculated in a time bin after the stimuli onset on each trial. Here we used as a time bin the first 100 ms following stimulus input onset; we found that major findings were qualitatively similar if later time bins are used (not shown). The SVM classifier was trained on half of the trials (randomly selected) and the other half of the trials were used for testing and prediction. The linear SVM classifier works by first constructing an optimal hyperplane for classification, based on labeled training data, and then generating predictions of the labels on testing data. Accuracy of the decoding is assessed by comparing the predicted labels to the actual labels. The accuracy is calculated as the proportion of trials classified correctly.

Specifically, the SVM receives TC activity as input: *r*^TC^(*i*_trial_, *i*_stim_, *i*_neu_, *i*_mod_) is the firing rate of the TC neuron *i*_neu_ with the *i*_stim_th level stimulus and *i*_mod_th level top-down inhibiton to RE neurons in the *i*_trial_ trials. Here *i*_trial_ = 1, …, 400, *i*_neu_ = 1, …, 1000, *i*_stim_ = 0, 1, …, 10, and *i*_mod_ = 1, …, 8. Thus the total dimension of *r*^TC^ is 400 × 11 × 1000 × 8.

For detection, the training set consists of 200 of the trials (across all TC neurons) with no stimulus (*i*_stim_ = 0 for each level of top-down modulation (with a dimension of 200 × 1 × 1000 × 8). The training set also consists of 20 trials for each level of stimulus *i*_stim_ = 1, …, 10 and each level of top-down modulation via (with a dimension of 20 × 10 × 1000 × 8). The training set thus includes half of the trials with no stimulus, and half with stimulus of various magnitudes. The testing set similarly consists of 200 trials with no stimulus (with a dimension of 200 × 1 × 1000 × 8, using trials that were not used for training), and 20 trials for each level of stimulus and top-down modulation (with a dimension of 20 × 10 × 1000 × 8, using trials that were not used for training). The labels for SVM for detection are presence (1) and absence (0) of stimulus.

For discrimination between two distinct structured inputs (input 1 and input 2) (within one modality), the training set includes 200 trials across all neurons for each level of stimulus strength (from 0 to 10) and top-down modulation via RE (from 1 to 8), for each of input 1 and 2. The dimension of the training set is thus 200 × 11 × 1000 × 8 × 2, where the ×2 accounts for input 1 and input 2. The trials not used for training have the same dimensions with the training set and is used for testing. The labels for SVM for discrimination are the identity of input 1 (1) and input 2 (0).

For congruent and incongruent discrimination across modalities, the training set includes 200 trials across levels of stimulus strength (from 0 to 10), using 2000 neurons (1000 TC neurons for each of the two modalities). Two conditions of top-down modulations are considered (for the model to attend to either modality and ignore the other) : *i*_mod_ = 8 for the first modality and *i*_mod_ = 1 for the second modality; and *i*_mod_ = 1 for the first modality and *i*_mod_ = 8 for the second modality.

The dimension of the training set is thus 200 × 11 × 2000 × 2. The testing set includes the other 200 trials not used in the training set, across the same conditions as for the training set. The labels for SVM for congruent and incongruent discrimination are the identity of input 1 (1) and input 2 (0) of the attended modality.

Detection and discrimination sensitivity (**Figures 2, S2, S3, and S5**) are defined as the slope of the detection and discrimination accuracy as a function of stimulus amplitude, within the linear portion with low stimulus amplitude.

### SVM, Stimulus, and Top-down Weight Vectors

The SVM generates a readout weight vector 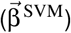, with which an SVM score (*S*_SVM_) can be calculated from the activity of TC neurons 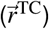:

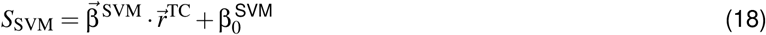

The activity of the *i*th TC neuron 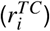 in the training data can be fitted to a linear regression model:

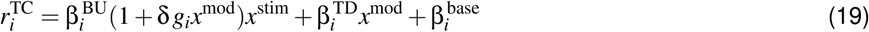

where *x*^stim^ is the dummy variable describing the strength of bottom-up stimulus input to TC neurons, *x*^mod^ is the dummy variable describing the strength of top-down input (inhibition to RE neurons in **Figure 4**.), and δ *g_i_* is the gain modulatory factor. 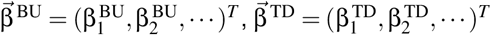, and 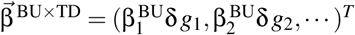 thus represent the regression coefficient weights across all neurons for bottom-up stimulus, top-down modulation, and their interaction.

The cosine similarities between the three regression weight vectors and 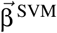 can be computed. In particular, we found that the SVM weight vector is positively correlated with 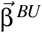 and 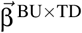, but is weakly negatively correlated with 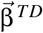 (**Figure 4D-E**). This can be understood by considering the SVM score, rewriting **Equation 18** as

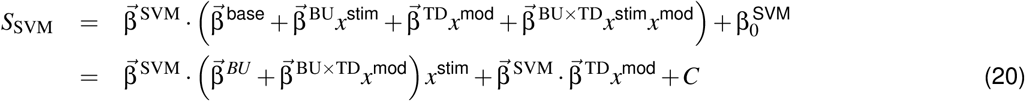

where 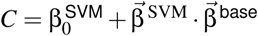 is a constant. Note that, the magnitude of each coefficient vector can be affected by the dummy variable. Thus, we consider their spatial orientation by calculating the cosine similarity, that is, studying the angle between each two vectors. The positive 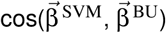 verifies that the SVM score is indeed sensitive to the presence of stimulus. Top-down modulation effectively increasing acting as a gain increase of the SVM sensitivity to bottom-up input (from 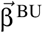 to 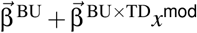). The negative 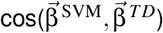 reduces the false alarm rate in the absence of stimulus (*x*^stim^ = 0), such that top-down modulation improves performance on stimulus-absent trials, by suppressing the SVM score and increasing the separation of SVM scores between trials with and without stimulus (**Figure 4**).

### Sensitivity Analysis

Perturbations to cellular and synaptic parameters were performed and their effect on multiple circuit and behavioral features were evaluated (**Figure S5**). The 6 perturbed parameters were (i) background excitatory input to TC neurons 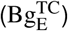, (ii) T-type calcium conductance in TC neurons 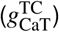, (iii) background excitatory input to RE neurons 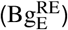, (iv) T-type calcium conductance in RE neurons 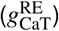, and (v, vi) the synaptic connections between the two neurons 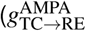 and 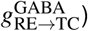. The 7 evaluated circuit and behavioral features were (i) TC firing rate (*r*_0_), (ii) response gain (*k*), (iii) spindle probability (*P*), (iv) detection sensitivity (*S*^det^), (v) discrimination sensitivity (*S*^disc^), (vi) congruent discrimination sensitivity (*S*^cong^), and (vii) incongruent discrimination sensitivity (*S*^inco^). In particular, the 6 perturbed parameters are varied by ±10%, and the feature sensitivity is defined as the change in the feature over the change in parameters. For instance, a feature sensitivity of 1 means that a 1% parameter perturbation alters the feature by 1%.

Specifically, we analyzed the feature sensitivity in the space of evaluated features, i.e., each perturbation is characterized by a column vector consisting of 7 different feature sensitivities. Therefore, we can obtain the feature sensitivity matrix *X* of dimension 7 × 6 within which each column represents one kind of perturbation (**Figure S5B**).

Singular value decomposition (SVD) is then performed on the sensitivity matrix *X* to find its principal components. The SVD of *X* is the factorization of *X* into the product of three matrices:

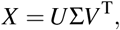

where *U* and *V* are 7 × 7 and 6 × 6 unitary matrices, respectively. Σ is a rectangular diagonal matrix with positive real entries (corresponding singular values, usually assuming σ_1_ ≥ σ_2_ ≥ …). Then, with the knowledge of linear algebra, one has

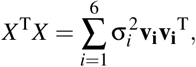

where **v_i_** is the *i*th column vector of *V*. Therefore, we define 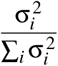, which represents the relative amount of reconstruction error reduced by including component *i*, as the fraction of the power in the feature sensitivity matrix associated with component *i*. It can be seen clearly that SVD is essentially principal component analysis (PCA) without mean subtraction. The first principal component from PCA captures the dominant pattern of variation. By contrast, the first component of SVD captures the dominant pattern of perturbation effects across features: **u_1_σ**_1_**v_1_**^T^ is the closest rank-1 approximation of *X*. In the SVD analysis, we found that SV1 explains 85% of the power in feature sensitivity, which indicates that the circuit’s sensitivity to perturbations is indeed approximately one-dimensional (**Figure S5C**). We found that the weights of SV1 are all the same sign, indicating that all features increase or decrease together along this axis (**Figure S5B** right). Finally, we examined the SV1 scores of the perturbation points, i.e. their projection along the SV1 axis. TC- and RE-related perturbations have opposite-sign scores, with synaptic perturbations at intermediate scores. We can therefore characterize the dominant mode of sensitivity as capturing alteration in excitation-inhibition balance (**Figure S5D**).

### Experimental Datasets of Neuronal Recordings

Full experimental details for the datasets have been previously described in Wimmer et al. (2015) and Nakajima et al. (2019). In brief, for this cross-modal sensory selection task, freely behaving mice were guided by a rule signal to select between spatially conflicting visual and auditory cues to receive a reward. At the beginning of each trial, mice were presented with 100 ms of either brown (10-kHz low-pass-filtered white noise) or blue noise (11-kHz high-pass-filtered white noise), each of which indicated one of two rules for sensory selection: attend to vision or attend to audition. This rule cue was followed by a delay period (0.4-0.6 s) prior to target stimulus presentation. In “attend to vision” trials (brown noise), the location of the rewarded port was signaled by a white LED (visual target) mounted underneath it in order to establish an association between the location of the visual target and the location of the reward port. In “attend to audition” trials (blue noise), mice learned the association between the auditory target, up-sweep (10-15kHz) or a down-sweep (16-12kHz) with right or left ports, respectively. The auditory and visual target stimuli were co-presented indicating reward at opposing locations and mice used rule cues (brown or blue noise) to match the target modality (vision vs. audition).

The dataset was recorded from two mice in the above cross-modal sensory selection task, including auditory thalamus (ventral medial geniculate body, MGBv), auditory thalamic reticular nucleus (audTRN), and primary auditory cortical (A1) excitatory and inhibitory neurons. As described in Nakajima et al. (2019), in order to make sure mice did not exhibit strong behavioral biases toward one modality and that they were able to use trial cues to choose modalities, only sessions with balanced performance (both auditory and visual trial types above 65%) across target modalities were used for the further analysis. Neurons were selected for analysis by filtering based only on data quality. We first excluded neurons which did not have any spikes during the stimulus presentation epoch over all the correct trials. The stimulus present epoch is defined from 20 ms after stimulus onset to the time of stimulus offset. We also excluded neurons which were not recorded stably throughout a condition during the stimulus presentation epoch. These were identified by calculating the coefficient of variation (CV) of each unit’s spike counts over trials of one stimulus at one modality condition. Units with CVs greater than 2 were deemed to be unstable and were excluded.

### Analysis of Neuronal Recordings

In the following analysis, we further selected MGBv neurons that have a significant preference to the up- or down-sweep stimulus. That is, these neurons exhibit stimulus tuning in their spiking activity which can support decoding of stimulus identity. The significant preference was determined by a two-tailed Mann-Whitney U test with α = 0.05. We then labeled a unit as preferring up-sweep (down-sweep) when it has a higher average firing rate to up-sweep (down-sweep) stimulus. 52 neurons satisfied this tuning criterion. We grouped the MGBv neurons with the four conditions (“preferred, attend to audition”, “nonpreferred, attend to audition”, “preferred, attend to vision”, and “nonpreferred, attend to vision”), and performed the analysis in **Figure 5**. In **Figure 5C**, we normalized the firing rate of each unit by the averaging firing rate of all correct trials in the pre-stimulus epoch (which is defined as total 100 ms just before the stimulus onset), and then smoothed the PSTH of each conditions with a causal exponential function with a 15-ms time constant. The normalized firing activities were aligned to the time of stimulus-onset.

For SVM-based decoding analysis on the MGBv neurons’ responses during the stimulus presentation epoch, similar to the decoding analysis in Nakajima et al. (2019), neurons were first pooled into a pseudo-population (**Figures 5E, S6A-C**). The SVM classifier was used for determine whether a firing activity pattern was evoked from the preferred stimulus or not. Note that, only correct trials were used and sessions were only included if greater than 20 correct trials per condition were available. In order to obtain the SVM accuracy, evoked spike counts under each condition were first randomly divided into training and test subsets (10 trials sampled 8 used for training and 2 for testing). Then according to the trials in the training subset, we labeled the responses of each unit as “preferred, attend to audition”, “nonpreferred, attend to audition”, “preferred, attend to vision”, and “nonpreferred, attend to vision”. In order to get a training firing activity pattern of a given condition, we randomly chose one evoked firing activity of each neuron from its pre-selected training subset under that condition and formed a firing activity vector. With a bootstrapping procedure, we generated 800 training firing activity vectors for each condition. In the same way, we generated 200 testing firing activity vectors for each condition. In the following, we ignored the attention rules (i.e., “preferred, attend to audition” + “preferred, attend to vision” → “preferred”, and same for “nonpreferred”), and trained the SVM classifier to distinguish the firing patterns only determined by “preferred” and “nonpreferred” conditions. The accuracy either in attention to audition trials or vision trials then could be obtained accordingly. To determine the variability of this estimate, the same procedure was repeated 100 times.

We used all the correct trials to determine the SVM axis of MGBv recordings (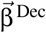 in **Figure 5G, I**). In the same way described by the above paragraph, we generated 1000 training firing activity vectors for each condition, and trained the SVM classifier of firing patterns ignoring the attention rules. In addition, we obtained the weights of 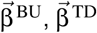 and 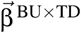 by fitting the linear regression model in **Equation** (19) with different dummy variables *x*^stim^ = 1 (0) for stim = “prefer” (“nonprefer”), and *x*^mod^ = 1 (−1) for mod = “attend to audition” (“attend to vision”). The results were robust to dummy variable design. In order to compare the MGBv recording analysis to our model results, we obtained the firing rate trials from the spiking circuit model within the detection paradigm (**Figure 2C**), only with stimuli of 0 and 0.4 μA/cm^2^ and with two levels of top-down modulation (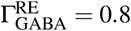 and 1.2), to be analogous to the experimental four conditions. Then we calculated all the coding axes discussed in our work (**Figure 5F–I**).

With the same selection rules above, we obtained 25 neurons in A1. Then, based on **Equation** (19), we calculated the gain modulation factor δ *g* with a 20-ms sliding window (**Figure 6C–D**). After selection of neurons, very few MGBv and A1 neurons were simultaneously recorded. Here, we predicted the PSTH of the *i*th neuron in the target A1 population by the PSTHs in the source MGBv population with a linear model:

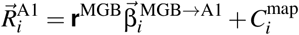

where 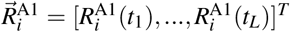 is a vector of firing activity at each given time point. The matrix **r**^MGB^ is *L* × *n* with *L* time points and *n* MGB neurons. Specifically, **r**^MGB^[*k*, *j*] represents the firing activity of the *j*th MGBv neuron at time point 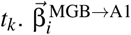 is the mapping vector from MGBv population to the *i*th A1 neuron. 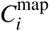 is a constant. To reduce overfitting, we used ridge regression with a regulation factor 2. We also considered the reconstruction with a time lag δ *t*, that is, **r**^MGB^[*k*, *j*] represents the firing activity of the *j*th MGBv neuron at time point *t_k_* + δ *t* (**Figure S6** D–E).

In addition, as shown in **Figure S6** F–G, we calculated the mapping vector 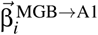 from the Poisson generalized linear model (GLM) (Keeley et al., 2020). In the Poisson GLM, the instantaneous firing rate of A1 approximated by the above equation with a nonlinear kernel:

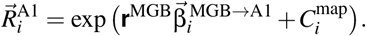

Then the spike count 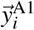 in the time bin of size Δ*t* (=20 ms, sliding window) follows a conditional Poisson distribution:

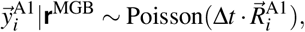

where the parameters were estimated by the maximum likelihood approach.

### Software Availability

Spiking circuit simulations were performed using the Brian2 simulator (Stimberg et al., 2019). Upon publication, custom modeling and analysis codes written in Python will be made publicly available on Github (https://github.com/murraylab), and a Brian2 implementation of the spiking circuit model will be made publicly available via the ModelDB model database (https://senselab.med.yale.edu/modeldb/).

## Supplementary Figures

**Figure S1:**
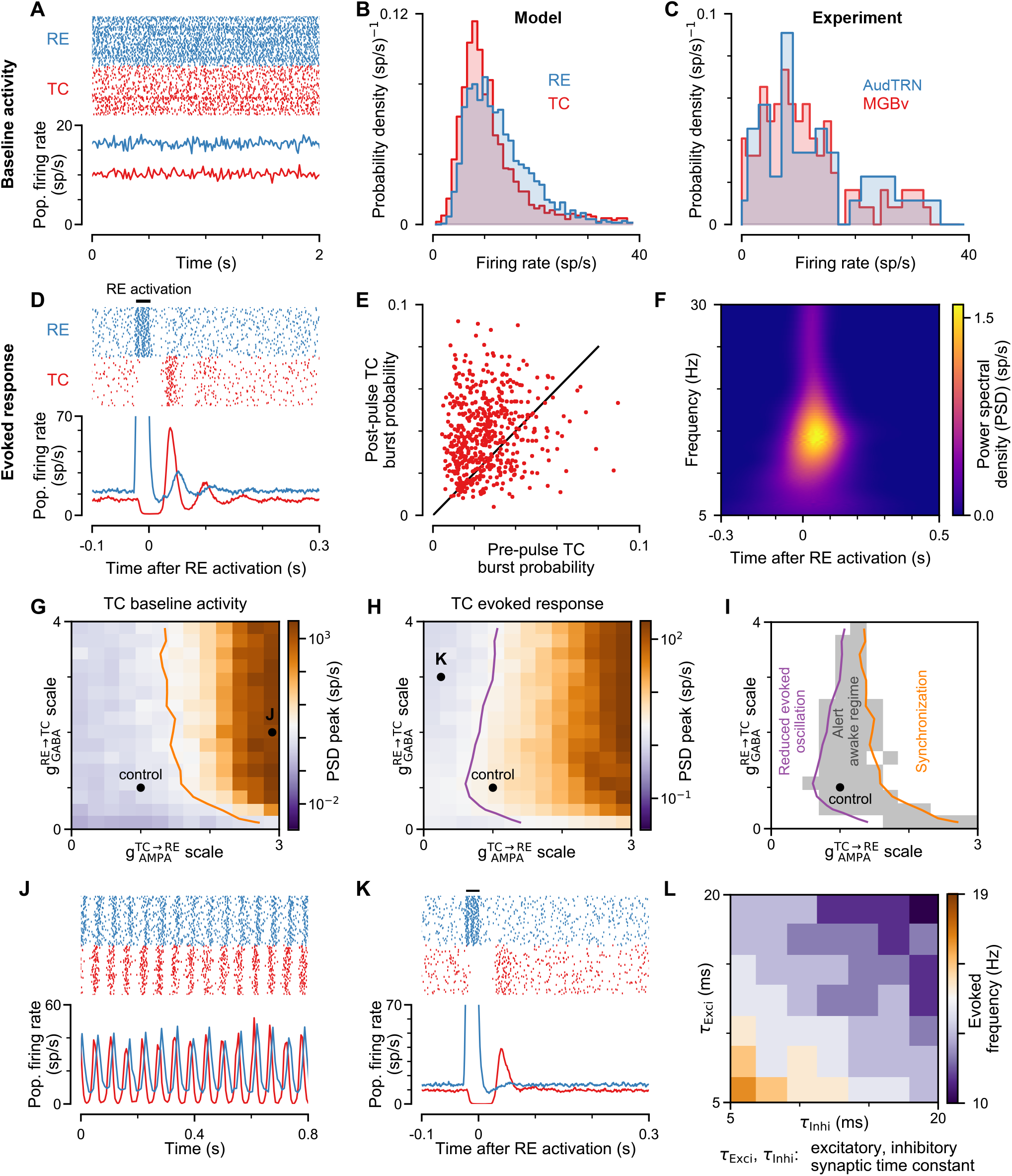
thalamic circuit model in alert awake regime. The circuit model is constrained to capture key circuit dynamics observed in alert behaving animals. **(A)** Rastergram showing quasi-asynchronous spiking activity in baseline state for RE (blue) and TC (red) neurons. **(B)** Distribution of TC and RE firing rates, which arises from heterogeneous recurrent and background connectivity. **(C)** Empirical distributions of firing rates in auditory thalamus (MGBv) and auditory thalamic reticular nucleus (AudTRN), recorded in awake behaving mice (Nakajima et al., 2019). The average firing rates of MGBv and auditory RE neurons are 11.2 sp/s and 13.8 sp/s, respectively (N=2 mice; n=119 MGBv; n=44 audTRN). Rastergram and evoked oscillation activity in response to pulse input to RE neurons. Black bar denotes time of pulse onset. **(E)** Burst probability post-pulse relative to pre-pulse. Burst probability is increased after RE pulse input. **(F)** (F) Power spectral density (PSD) over time of evoked response to RE pulse. The evoked response demonstrates decaying oscillation in the spindle frequency. **(G)** Peak value of the PSD for spontaneous activity, as a function of 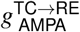 and 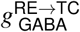. Oscillation arises from strong coupling from TC to RE 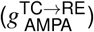. Orange line denotes the upper bound where the circuit exhibits quasi-asynchronous spontaneous activity (see **Methods**). **(H)** PSD peak for the evoked response to RE pulse input as a function of 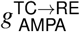 and 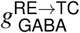. Spindle oscillation is reduced with weak coupling from TC to RE 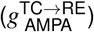 or strong coupling from RE to TC 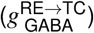. Purple line denotes the lower bound where the circuit exhibits an evoked spindle-like response consistent to experimental observations. **(I)** Parameter ranges of 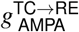 and 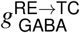 considered within the alert awake regime (shaded). Black dot denotes the control circuit. **(J)** Due to strong synaptic connectivity between TC and RE populations, the circuit model exhibits strong spontaneous synchronous oscillations. Parameters are marked in **G**. **(K)** The synaptic conductances of the circuit model are too weak to generate reverberating oscillations when perturbed. Parameters are marked in **H**. **(L)** Evoked oscillation frequency of the circuit model with different synaptic time constants. Here, to preserve total synaptic strength, conductances are scaled inversely with the scaling of time constants. These results indicate that the intrinsic oscillation frequency in the circuit is shaped by both excitatory and inhibitory synaptic time constants.

**Figure S2:**
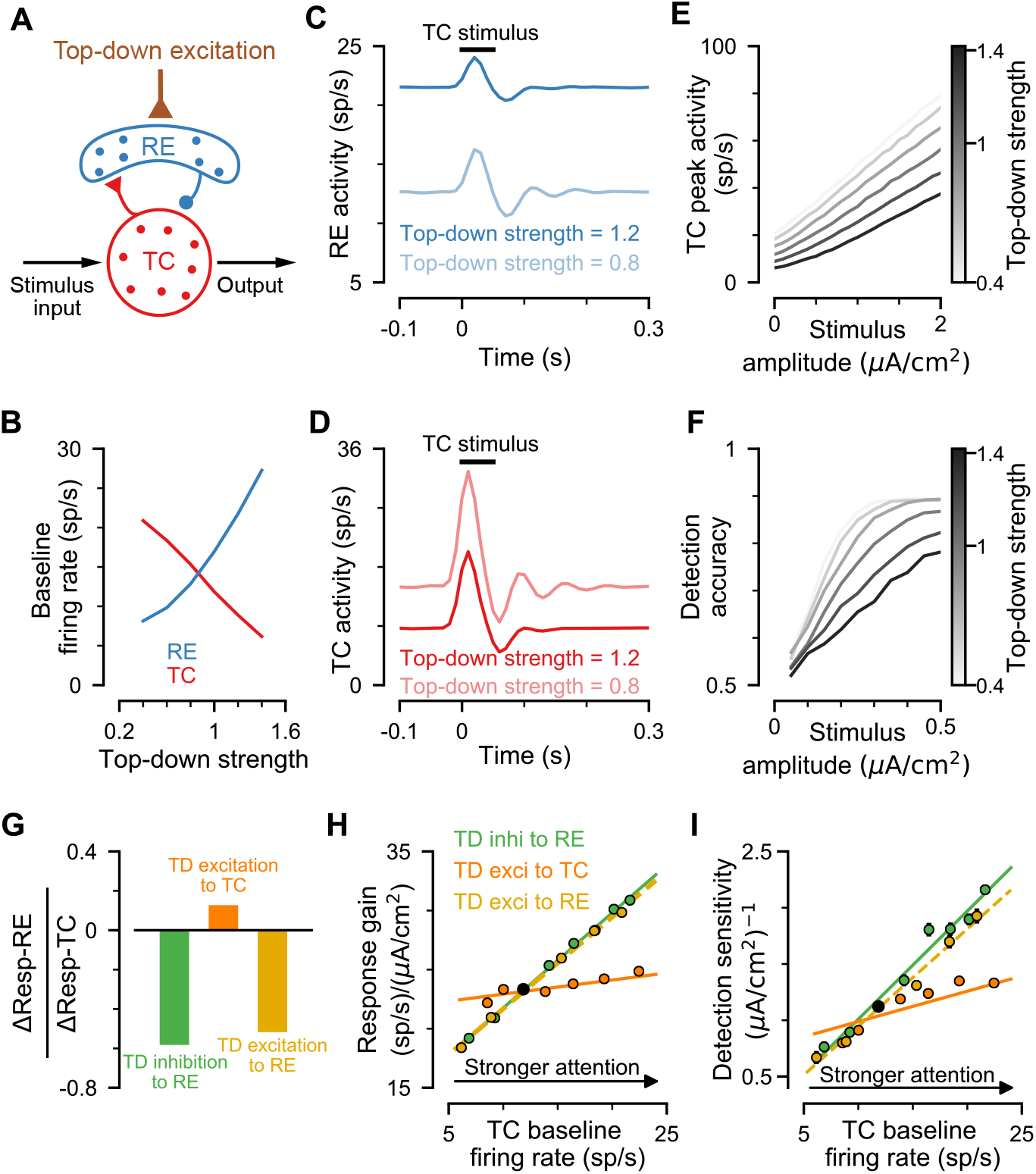
Top-down Excitation to TC or to RE. **(A)** Schematic of top-down excitation to RE neurons. **(B)** RE (blue) and TC (red) baseline firing rate as a function of top-down modulation strength (excitation to RE). Of note, the direction of modulation here is opposite of that for inhibition to RE **Figure 2**. **(C)** RE population activity in response to stimulus input with different top-down modulation strengths. Black bar denotes the time of TC stimulus onset. **(D)** TC Population activity in response to stimulus input with different top-down modulation strengths. Black bar denotes the time of TC stimulus onset. Gain curves of TC sensory outputs under different levels of top-down modulation strength. Darker color represents stronger top-down strength. **(F)** Detection accuracy under different levels of top-down modulation strength. Darker color represents stronger top-down strength. **(G)** The ratio firing-rate changes for RE and TC neurons, for top-down modulation scenarios: inhibition to RE, excitation to TC, excitation to RE. The same change in TC firing rate can be obtained from the three different top-down modulations, but RE responses differentiate TD→TC from TD→RE scenarios. **(H)** Response gain of the circuit, as a function of top-down modulation strength, for the three scenarios. **(I)** Sensitivity of detection accuracy as a function of top-down modulation strength. Panels **(H,I)** show that top-down modulation via RE neurons has a stronger effect on response gain and detection performance, than via TC neurons. Black dots represent the control parameter set.

**Figure S3:**
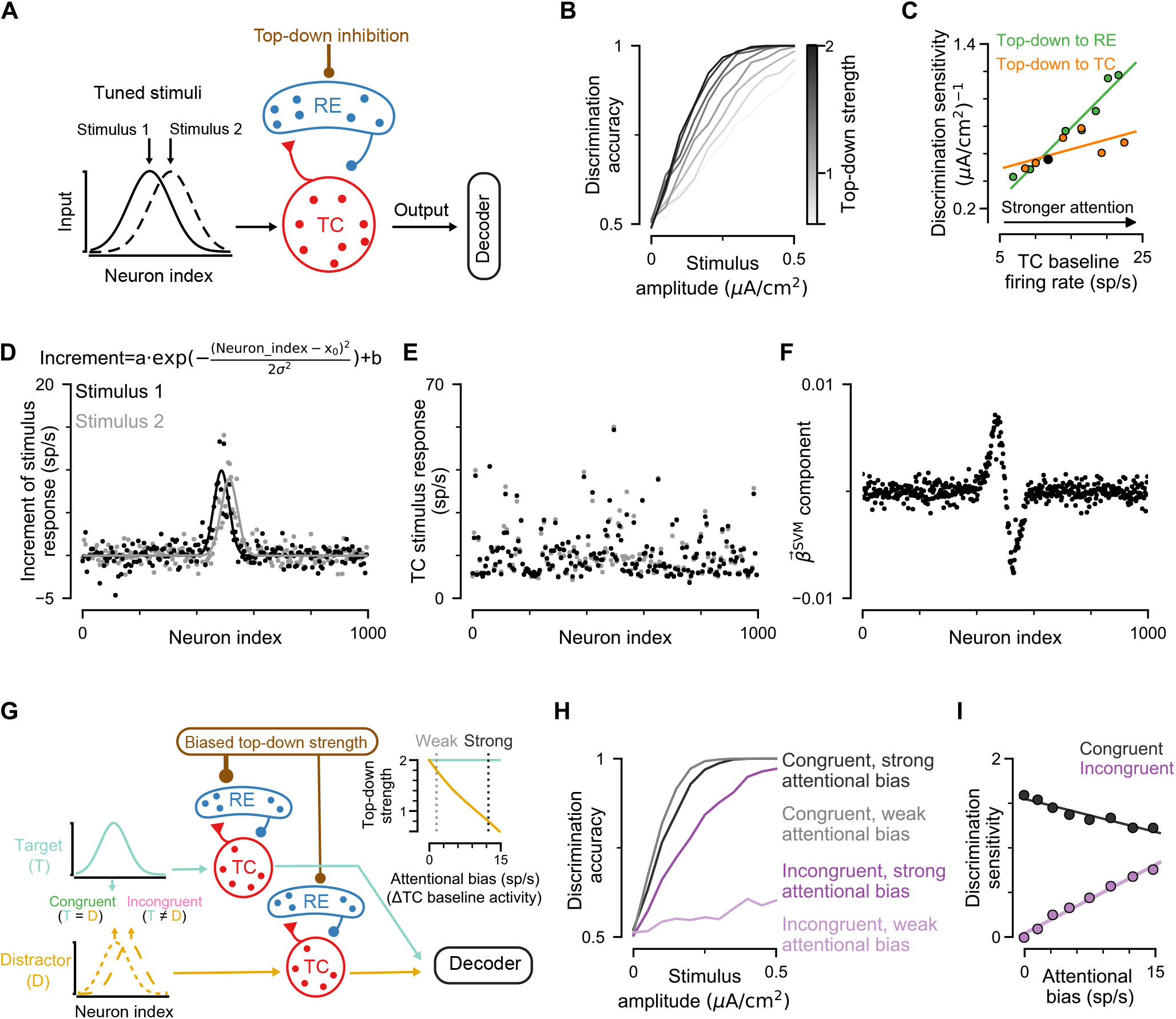
Attentional control of performance in discrimination paradigms. **(A)** Schematic of uni-modal discrimination paradigm. Two tuned, partially overlapping stimulus patterns are used as inputs to the TC population. On each trial, one stimulus is presented, with variable amplitude to control difficulty. A downstream decoder is trained on TC output to classify which stimulus was presented. Top-down modulation is via non-specific inhibition to RE. **(B)** Discrimination accuracy of a downstream decoder under different levels of top-down modulation strength. Darker colors represent stronger top-down modulation. **(C)** Discrimination sensitivity of discrimination accuracy as top-down strength is varied, for modulation via RE vs. via TC. **(D)** The TC firing rate with the tuned input as a function of the neuron index after subtracting the baseline firing rate. This shows the tuning and overlap of stimulus input patterns. The solid lines are fitted with top equations (Gaussian profile). **(E)** The TC firing rate as a function of the neuron index with the tuned input. The firing rate does not clearly show any tuned profile. **(F)** The component profile of SVM weight vector as a function of the neuron index. It shows that there are distinct peaks at the centers of the tuned Stimulus 1 (positive) and Stimulus 2 (negative). **(G)** Schematic of the cross-modal (e.g., visual and auditory) discrimination paradigm. Here there are two TC-RE circuits, each corresponding to a different modality. Biased top-down modulation via RE across modalities allows attention toward the target modality (cyan) and filtering of the distractor modality (yellow), as well as contextual switching of which modality is target vs. distractor. The target and distractor stimuli can be either congruent, such that they map to the same readout response, or incongruent, such that they provide contrasting information. (Inset) Top-down strengths target and distractor modalities as a function of the attentional bias. The attentional bias is defined as the difference between the TC baseline activity in the attended modality to that in unattended modality (i.e., ΔTC baseline activity). **(H)** Discrimination accuracy of a downstream decoder, for congruent (black) and incongruent (purple) trials with strong (dark) and weak (light) attentional bias. **(I)** Discrimination sensitivity as a function of attentional bias, for congruent and incongruent trials. Attentional bias can enhance discrimination accuracy for incongruent trials by filtering out distractor stimuli. Across all conditions, the top-down modulatory input into the attended modality is held constant.

**Figure S4:**
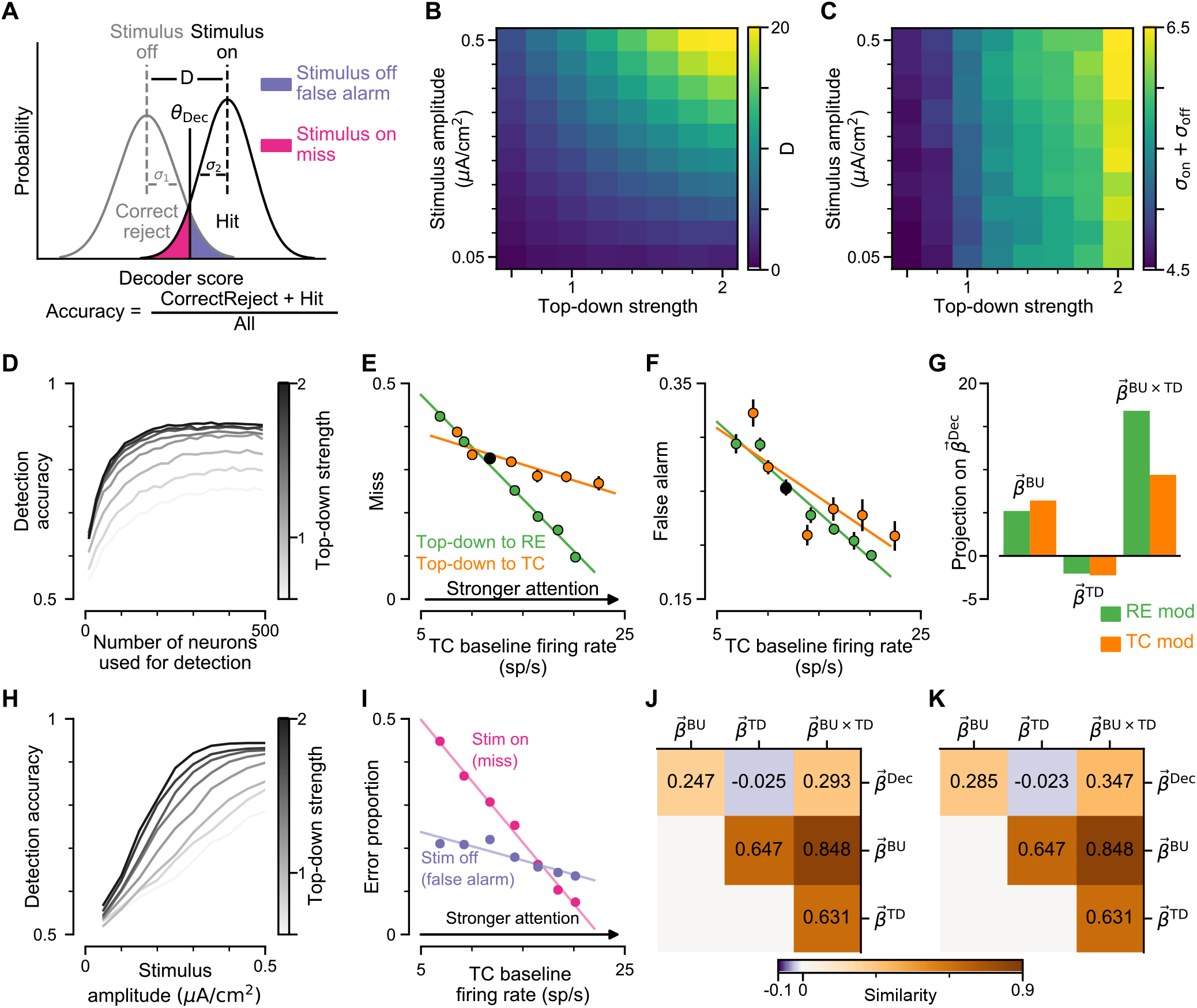
SVM decoder for detection and effects of top-down modulation via RE and TC neurons on decoding for detection. **(A)** Schematic of signal detection with the SVM decoder. Detection accuracy depends on the difference (D) between the mean of the SVM score distributions of stimulus present and absent, as well as, their standard deviations (σ_1_, σ_2_). **(B)** The mean difference between SVM score distributions of stimulus on and off as a function of bottom-up stimulus strength and top-down strength to RE. Both factors increase the score mean difference (*D*) with a strong impact. (C) The sum of standard deviations (σ_on_ + σ_off_) as a function of bottom-up stimulus and top-down strength to RE. Both factors have small effect on the deviation (note small range). These results indicate that stimulus and top-down inputs improve detection performance through their effects on the score difference *D*. **(D)** Detection accuracy as a function of the number of neurons used in the SVM decoder. Proportion of stimulus-on trials which are misses, as a function of top-down modulation strength, with top-down modulation via either inhibition to RE neurons (green) or via excitation to TC neurons (orange). Here, TC baseline activity denotes a common proxy for the strength top-down modulation, to compare RE- and TC-mediated modulations. The rate of misses is reduced more effectively by RE modulation than by TC modulation. **(F)** Proportion of stimulus-off trials which are false alarms, as a function of top-down modulation strength, with top-down modulation via either inhibition to RE neurons (green) or via excitation to TC neurons (orange). The rate of false alarms is reduced similarly by both modulations. **(G)** Projections of the bottom-up stimulus, top-down modulation, and interaction weight vectors on the SVM vector weight. Top-down modulation via RE neurons (green) has a larger interaction term than top-down modulation via TC neurons, which is consistent with the results shown in **Figure 2I** (i.e., RE modulation has a larger effect). The top-down modulation of baseline activity helps to reduce the false alarm rate, and top-down effect on output gain helps to reduce the miss rate. **(H)** Detection accuracy as a function of the stimulus amplitude. Here, the SVM decoder was trained at a single level of top-down modulation strength (1.4). **(I)** Error proportion as a function of top-down strength. Both miss and false alarm trials are reduced in the SVM decoder of **(H)**. **(J,K)** Cosine similarity between different weight vectors using an SVM trained at a single level of top-down modulation strength **(J)**, and using an SVM trained at multiple levels of top-down modulation strength **(K)**. Both **(J)** and **(K)** consistently show that there are the positive similarities between the bottom-up, top-down and interactions, whereas the decoder axis is negatively correlated with the top-down vector. This negative projection helps the decoder to distinguish between top-down attentional inputs or bottom-up stimulus inputs, even though both increase TC firing rates overall.

**Figure S5:**
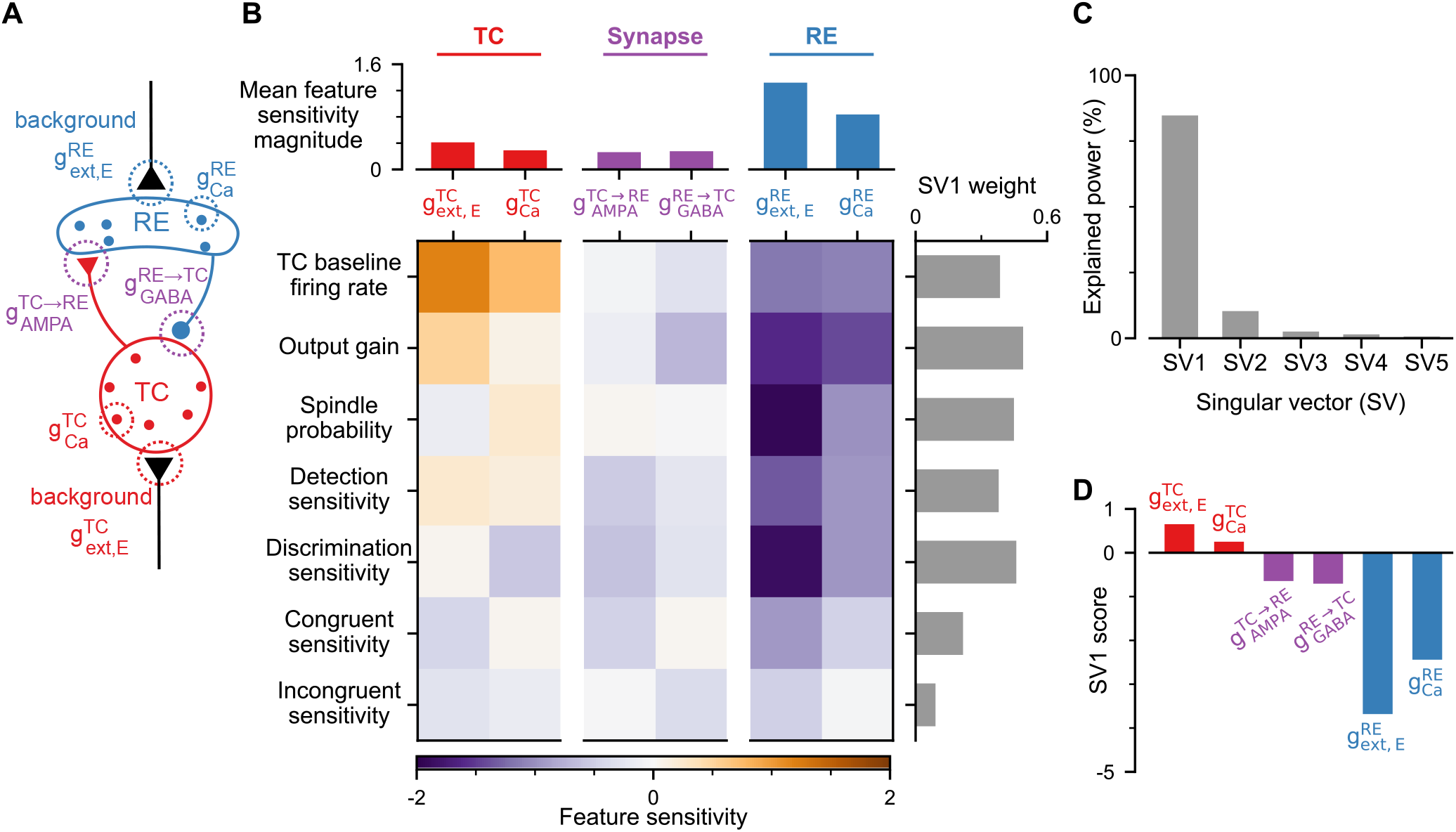
Sensitivity analysis of circuit dynamics and computations. **(A)** Schematic of biophysical parameters subject to perturbation, related to TC (red), RE (blue), and synaptic (purple) properties. Six biophysical parameters (4 synaptic and 2 cellular) were background excitatory AMPA conductances (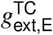 and 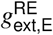), cellular calcium conductances (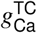 and 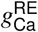) and the synaptic connections between TC and RE populations (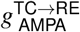 and 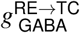). **(B)** (Bottom) Sensitivity value of seven dynamical and computational features to various perturbations. Feature selectivity is defined as the ratio of the proportional change in the feature to the proportional change in the parameter. A feature sensitivity of 1 means a 1% perturbation would drive a 1% change in the feature. (Top) Mean feature sensitivity amplitude across features for each perturbation. Averaging across all features, the RE-specific parameters generated greater feature sensitivities than did TC-specific or synaptic parameters. This finding is consistent with the conclusion that RE neurons are a potent site for modulation of thalamic processing, as well as with proposals that RE neurons may be a site of vulnerability. **(B)** (Right) Weights of the first singular vector (SV1) of the feature selectivity matrix, in terms of the feature components, derived from singular value decomposition (SVD). SVD is essentially principal component analysis but without mean subtraction (see **Methods**). The singular vectors from SVD associated with the top *k* singular values define the closest approximation to the feature sensitivity matrix by a rank-*k* matrix. Considering each of the six perturbations as a point in the seven-dimensional space of features, SV1, the leading singular vector, defines the axis in feature space that maximally captures the patterns of feature alterations across perturbations. **(C)** Explained power of the leading singular vectors. SV1 explains 85% of the power in feature sensitivity matrix across perturbations, revealing a quasi-one-dimensional effect of biophysical perturbations on the set of features. In the right panel of **(B)**, the weights of SV1 are all the same sign, indicating that all features increase or decrease together along this axis. **(D)** SV1 scores of the perturbations (i.e. projections of the perturbation vector along SV1). TC perturbations vs. RE perturbations are separated, revealing the quasi-one-dimensional dependence of the circuit on perturbation to excitation-inhibition balance.

**Figure S6:**
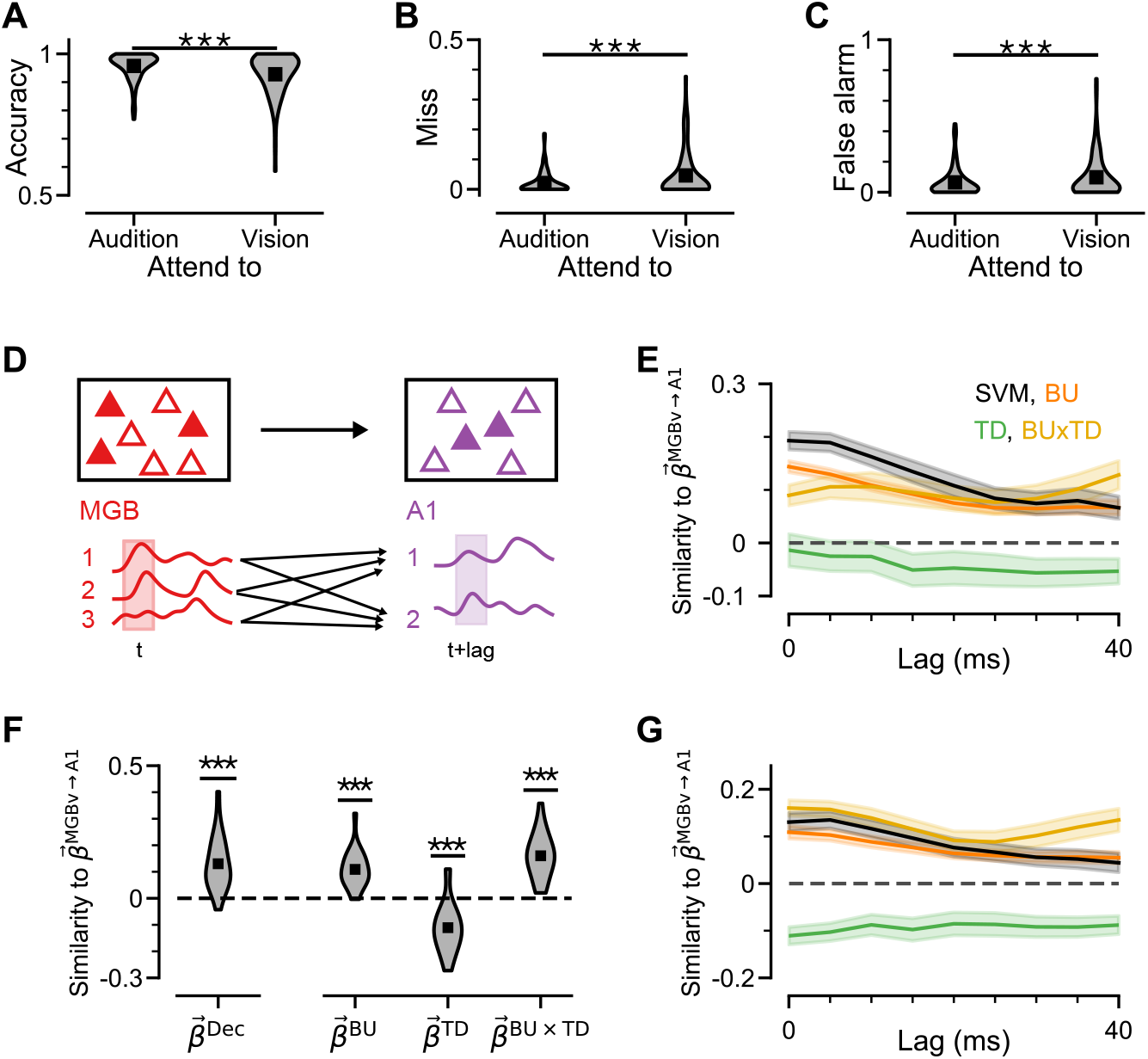
Analysis on MGBv and A1 recordings. **(A)** SVM decoder based on MGBv neuronal firing activities evoked by their preferred and nonpreferred stimuli. The SVM here was trained by trials from both “attend to audition” and “attend to vision” conditions. Target sounds were more easily classified during “attend to audition” trials compared with “attend to vision” trials (Wilcoxon test, *p* < 0.001; N=2 mice, n=52 neurons). **(B,C)** Miss likelihood (when stimulus present) and false alarm likelihood (when stimulus absent) of the SVM classifier. Both misses and false alarms were reduced in the “attend to audition” condition, relative to “attend to vision” (Wilcoxon test, *p* < 0.001). **(D)** Schematic of how the peri-stimulus-time histograms (PSTHs) of target A1 neurons are related to MGBv neurons, here fit with a time lag between areas (see **Methods**). **(E)** Cosine similarity between mapping weights from MGBv to A1 neurons and four coding axes, across a range of time lags. The relationships were robust, in that the signs (positive/negative) of similarities remained consistent across different time lags. **(F)** Cosine similarity between mapping weights from MGBv to A1 neurons and coding axes obtained in **Figure 5**. Here, different from **Figure 6F**, the mapping weights were calculated with a Poisson generalized linear model (GLM) (see **Methods**). Consistent with the linear regression results in **Figure 6F**, here we found that mapping weights have a positive similarity with the decoder axis. Mapping weights have a positive, negative, and positive similarity with bottom-up, top-down, and interaction axis, respectively (Mann-Whitney U test, *p* < 0.001). **(G)** Same as in **(E)** but here using the Poisson GLM method. The relationships are robust in keeping the same sign when using the Poisson GLM method.

## References

Abbott L.F., and Chance F.S. (2005). Drivers and modulators from push-pull and balanced synaptic input. Prog Brain Res 149, 147–55.

Ahrens S., Jaramillo S., Yu K., Ghosh S., Hwang G.R., Paik R., Lai C., He M., Huang Z.J., and Li B. (2015). ErbB4 regulation of a thalamic reticular nucleus circuit for sensory sselection. Nat Neurosci 18, 104–11.

Bastos A.M., Briggs F., Alitto H.J., Mangun G.R., and Usrey W.M. (2014). Simultaneous recordings from the primary visual cortex and lateral geniculate nucleus reveal rhythmic interactions and a cortical source for gamma-band oscillations. Journal of Neuroscience 34, 7639–7644.

Bazhenov M., Timofeev I., Steriade M., and Sejnowski T. (2000). Spiking-bursting activity in the thalamic reticular nucleus initiates sequences of spindle oscillations in thalamic networks. J Neurophysiol 84, 1076–87.

Bazhenov M., Timofeev I., Steriade M., and Sejnowski T.J. (2002). Model of thalamocortical slow-wave sleep oscillations and transitions to activated states. The Journal of Neuroscience 22, 8691–8704.

Béhuret S., Deleuze C., and Bal T. (2015). Corticothalamic synaptic noise as a mechanism for selective attention in thalamic neurons. Frontiers in Neural Circuits 9.

Briggs F., and Usrey W.M. (2008). Emerging views of corticothalamic function. Curr Opin Neurobiol 18, 403–7.

Bruno R.M. (2006). Cortex is driven by weak but synchronously active thalamocortical synapses. Science 312, 1622–1627.

Clemente-Perez A., Makinson S.R., Higashikubo B., Brovarney S., Cho F.S., Urry A., Holden S.S., Wimer M., Dávid C., Fenno L.E., Acsády L., Deisseroth K., and Paz J.T. (2017). Distinct thalamic reticular cell types differentially modulate normal and pathological cortical rhythms. Cell Reports 19, 2130–2142.

Crandall S.R., Cruikshank S.J., and Connors B.W. (2015). A corticothalamic switch: controlling the thalamus with dynamic synapses. Neuron 86, 768–82.

Crick F. (1984). Function of the thalamic reticular complex: the searchlight hypothesis. Proc Natl Acad Sci U S A 81, 4586–90.

Destexhe A., Babloyantz A., and Sejnowski T.J. (1993a). Ionic mechanisms for intrinsic slow oscillations in thalamic relay neurons. Biophys J 65, 1538–52.

Destexhe A., Bal T., McCormick D.A., and Sejnowski T.J. (1996). Ionic mechanisms underlying synchronized oscillations and propagating waves in a model of ferret thalamic slices. J Neurophysiol 76, 2049–70.

Destexhe A., Contreras D., Sejnowski T.J., and Steriade M. (1994). A model of spindle rhythmicity in the isolated thalamic reticular nucleus. J Neurophysiol 72, 803–18.

Destexhe A., Contreras D., and Steriade M. (1998). Mechanisms underlying the synchronizing action of corticothalamic feedback through inhibition of thalamic relay cells. J Neurophysiol 79, 999–1016.

Destexhe A., McCormick D.A., and Sejnowski T.J. (1993b). A model for 8-10 Hz spindling in interconnected thalamic relay and reticularis neurons. Biophys J 65, 2473–7.

Fagerberg L., Hallström B.M., Oksvold P., Kampf C., Djureinovic D., Odeberg J., Habuka M., Tahmasebpoor S., Danielsson A., Edlund K., Asplund A., Sjöstedt E., Lundberg E., Szigyarto C.A.K., Skogs M., Takanen J.O., Berling H., Tegel H., Mulder J., Nilsson P., Schwenk J.M., Lindskog C., Danielsson F., Mardinoglu A., Sivertsson A., von Feilitzen K., Forsberg M., Zwahlen M., Olsson I., Navani S., Huss M., Nielsen J., Ponten F., and Uhlén M. (2014). Analysis of the human tissue-specific expression by genome-wide integration of transcriptomics and antibody-based proteomics. Mol Cell Proteomics 13, 397–406.

Ferguson K.A., and Cardin J.A. (2020). Mechanisms underlying gain modulation in the cortex. Nat Rev Neurosci 21, 80–92.

Fu Y., Tucciarone J.M., Espinosa J.S., Sheng N., Darcy D.P., Nicoll R.A., Huang Z.J., and Stryker M.P. (2014). A cortical circuit for gain control by behavioral state. Cell 156, 1139–1152.

Golomb D., Wang X.J., and Rinzel J. (1996). Propagation of spindle waves in a thalamic slice model. J Neurophysiol 75, 750–69.

Halassa M.M., and Kastner S. (2017). Thalamic functions in distributed cognitive control. Nat Neurosci 20, 1669–1679.

Halassa M.M., Siegle J.H., Ritt J.T., Ting J.T., Feng G., and Moore C.I. (2011). Selective optical drive of thalamic reticular nucleus generates thalamic bursts and cortical spindles. Nat Neurosci 14, 1118–20.

Hirai D., Nakamura K.C., ichi Shibata K., Tanaka T., Hioki H., Kaneko T., and Furuta T. (2017). Shaping somatosensory responses in awake rats: cortical modulation of thalamic neurons. Brain Structure and Function 223, 851–872.

Hou G., Smith A.G., and Zhang Z.W. (2016). Lack of intrinsic GABAergic connections in the thalamic reticular nucleus of the mouse. J Neurosci 36, 7246–52.

Huguenard J.R., and McCormick D.A. (2007). Thalamic synchrony and dynamic regulation of global forebrain oscillations. Trends in Neurosciences 30, 350–356.

Jaramillo J., Mejias J.F., and Wang X.J. (2019). Engagement of pulvino-cortical feedforward and feedback pathways in cognitive computations. Neuron 101, 321–336.e9.

Jones E.G. (2012). The thalamus (Springer Science & Business Media).

Keeley S.L., Zoltowski D.M., Aoi M.C., and Pillow J.W. (2020). Modeling statistical dependencies in multi-region spike train data. Current Opinion in Neurobiology 65, 194–202.

Kepecs A., and Fishell G. (2014). Interneuron cell types are fit to function. Nature 505, 318–26.

Krauzlis R.J., Bogadhi A.R., Herman J.P., and Bollimunta A. (2018). Selective attention without a neocortex. Cortex 102, 161–175.

Krishnan G.P., Chauvette S., Shamie I., Soltani S., Timofeev I., Cash S.S., Halgren E., and Bazhenov M. (2016). Cellular and neurochemical basis of sleep stages in the thalamocortical network. eLife 5.

Krol A., Wimmer R.D., Halassa M.M., and Feng G. (2018). Thalamic reticular dysfunction as a circuit endophenotype in neurodevelopmental disorders. Neuron 98, 282–295.

Li Y., Lopez-Huerta V.G., Adiconis X., Levandowski K., Choi S., Simmons S.K., Arias-Garcia M.A., Guo B., Yao A.Y., Blosser T.R., Wimmer R.D., Aida T., Atamian A., Naik T., Sun X., Bi D., Malhotra D., Hession C.C., Shema R., Gomes M., Li T., Hwang E., Krol A., Kowalczyk M., Peça J., Pan G., Halassa M.M., Levin J.Z., Fu Z., and Feng G. (2020). Distinct subnetworks of the thalamic reticular nucleus. Nature 583, 819–824.

Litwin-Kumar A., Rosenbaum R., and Doiron B. (2016). Inhibitory stabilization and visual coding in cortical circuits with multiple interneuron subtypes. Journal of Neurophysiology 115, 1399–1409.

Ly C., and Doiron B. (2009). Divisive gain modulation with dynamic stimuli in integrate-and-fire neurons. PLoS Computational Biology 5, e1000365.

Martinez-Garcia R.I., Voelcker B., Zaltsman J.B., Patrick S.L., Stevens T.R., Connors B.W., and Cruikshank S.J. (2020). Two dynamically distinct circuits drive inhibition in the sensory thalamus. Nature 583, 813–818.

McAlonan K., Cavanaugh J., and Wurtz R.H. (2008). Guarding the gateway to cortex with attention in visual thalamus. Nature 456, 391–4.

Mehaffey W.H. (2005). Deterministic multiplicative gain control with active dendrites. Journal of Neuroscience 25, 9968–9977.

Murray J.D., and Anticevic A. (2017). Toward understanding thalamocortical dysfunction in schizophrenia through computational models of neural circuit dynamics. Schizophr Res 180, 70–77.

Murray J.D., Jaramillo J., and Wang X.J. (2017). Working memory and decision-making in a frontoparietal circuit model. J Neurosci 37, 12167–12186.

Nakajima M., and Halassa M.M. (2017). Thalamic control of functional cortical connectivity. Curr Opin Neurobiol 44, 127–131.

Nakajima M., Schmitt L.I., and Halassa M.M. (2019). Prefrontal cortex regulates sensory filtering through a basal ganglia-to-thalamus pathway. Neuron 103, 445–458.e10.

O’Connor D.H., Fukui M.M., Pinsk M.A., and Kastner S. (2002). Attention modulates responses in the human lateral geniculate nucleus. Nat Neurosci 5, 1203–9.

Ozeki H., Finn I.M., Schaffer E.S., Miller K.D., and Ferster D. (2009). Inhibitory stabilization of the cortical network underlies visual surround suppression. Neuron 62, 578–92.

Phillips J.W., Schulmann A., Hara E., Winnubst J., Liu C., Valakh V., Wang L., Shields B.C., Korff W., Chandrashekar J., Lemire A.L., Mensh B., Dudman J.T., Nelson S.B., and Hantman A.W. (2019). A repeated molecular architecture across thalamic pathways. Nat Neurosci 22, 1925–1935.

Ramcharan E.J., Gnadt J.W., and Sherman S.M. (2005). Higher-order thalamic relays burst more than first-order relays. Proc Natl Acad Sci U S A 102, 12236–41.

Richard E.A., Khlestova E., Nanu R., and Lisman J.E. (2017). Potential synergistic action of 19 schizophrenia risk genes in the thalamus. Schizophr Res 180, 64–69.

Rikhye R.V., Wimmer R.D., and Halassa M.M. (2018). Toward an integrative theory of thalamic function. Annu Rev Neurosci 41, 163–183.

Ritter-Makinson S., Clemente-Perez A., Higashikubo B., Cho F.S., Holden S.S., Bennett E., Chkhaidze A., Rooda O.H.E., Cornet M.C., Hoebeek F.E., Yamakawa K., Cilio M.R., Delord B., and Paz J.T. (2019). Augmented reticular thalamic bursting and seizures in Scn1a-Dravet Syndrome. Cell Reports 26, 54–64.e6.

Roberts J.A., and Robinson P.A. (2012). Corticothalamic dynamics: structure of parameter space, spectra, instabilities, and reduced model. Phys Rev E Stat Nonlin Soft Matter Phys 85, 011910.

Saalmann Y.B., and Kastner S. (2011). Cognitive and perceptual functions of the visual thalamus. Neuron 71, 209–223.

Saleem A.B., Lien A.D., Krumin M., Haider B., Rosón M.R., Ayaz A., Reinhold K., Busse L., Carandini M., and Harris K.D. (2017). Subcortical source and modulation of the narrowband gamma oscillation in mouse visual cortex. Neuron 93, 315–322.

Salinas E., and Thier P. (2000). Gain modulation: a major computational principle of the central nervous system. Neuron 27, 15–21.

Sanzeni A., Akitake B., Goldbach H.C., Leedy C.E., Brunel N., and Histed M.H. (2020). Inhibition stabilization is a widespread property of cortical networks. eLife 9.

Schmitt L.I., Wimmer R.D., Nakajima M., Happ M., Mofakham S., and Halassa M.M. (2017). Thalamic amplification of cortical connectivity sustains attentional control. Nature 545, 219–223.

Semedo J.D., Gokcen E., Machens C.K., Kohn A., and Yu B.M. (2020). Statistical methods for dissecting interactions between brain areas. Current Opinion in Neurobiology 65, 59–69.

Semedo J.D., Zandvakili A., Machens C.K., Yu B.M., and Kohn A. (2019). Cortical areas interact through a communication subspace. Neuron 102, 249–259.e4.

Sherman S.M. (2012). Thalamocortical interactions. Current Opinion in Neurobiology 22, 575–579.

Sherman S.M. (2016). Thalamus plays a central role in ongoing cortical functioning. Nature neuroscience 19, 533–541.

Sohal V.S., Pangratz-Fuehrer S., Rudolph U., and Huguenard J.R. (2006). Intrinsic and synaptic dynamics interact to generate emergent patterns of rhythmic bursting in thalamocortical neurons. J Neurosci 26, 4247–55.

Stimberg M., Brette R., and Goodman D.F. (2019). Brian 2, an intuitive and efficient neural simulator. Elife 8.

Temereanca S., Brown E.N., and Simons D.J. (2008). Rapid changes in thalamic firing synchrony during repetitive whisker stimulation. Journal of Neuroscience 28, 11153–11164.

Timofeev I., Bonjean M.E., and Bazhenov M. (2020). Cellular Mechanisms of Thalamocortical Oscillations in the Sleeping Brain (New York, NY: Springer New York), pp. 119–170.

Usrey W.M., Reppas J.B., and Reid R.C. (1998). Paired-spike interactions and synaptic efficacy of retinal inputs to the thalamus. Nature 395, 384–7.

Wang X.J., Golomb D., and Rinzel J. (1995). Emergent spindle oscillations and intermittent burst firing in a thalamic model: specific neuronal mechanisms. Proc Natl Acad Sci U S A 92, 5577–81.

Wei H., Bonjean M., Petry H.M., Sejnowski T.J., and Bickford M.E. (2011). Thalamic burst firing propensity: a comparison of the dorsal lateral geniculate and pulvinar nuclei in the tree shrew. J Neurosci 31, 17287–99.

Wells M.F., Wimmer R.D., Schmitt L.I., Feng G., and Halassa M.M. (2016). Thalamic reticular impairment underlies attention deficit in Ptchd1^*Y* /−^ mice. Nature 532, 58–63.

Wimmer R.D., Schmitt L.I., Davidson T.J., Nakajima M., Deisseroth K., and Halassa M.M. (2015). Thalamic control of sensory selection in divided attention. Nature 526, 705–9.

Wong K.F., and Wang X.J. (2006). A recurrent network mechanism of time integration in perceptual decisions. J Neurosci 26, 1314–28.

Yang G.R., Murray J.D., and Wang X.J. (2016). A dendritic disinhibitory circuit mechanism for pathway-specific gating. Nat Commun 7, 12815.

## References

Jones E.G. (2012). The thalamus (Springer Science & Business Media).

Kepecs A., and Fishell G. (2014). Interneuron cell types are fit to function. Nature 505, 318–26.

Sherman S.M. (2012). Thalamocortical interactions. Current Opinion in Neurobiology 22, 575–579.

